# Uncovering the universality of self-replication in protein aggregation and its link to disease

**DOI:** 10.1101/2022.06.08.495339

**Authors:** Georg Meisl, Catherine K Xu, Jonathan D Taylor, Thomas C T Michaels, Aviad Levin, Daniel Otzen, David Klenerman, Steve Matthews, Sara Linse, Maria Andreasen, Tuomas P J Knowles

## Abstract

Fibrillar protein aggregates are a hallmark of the pathology of a range of human disorders, from prion diseases to dementias. Yet, the same aggregated structures that are formed in disease are also encountered in several functional contexts. The fundamental properties that determine whether these protein assembly processes are functional or, by contrast, pathological, have remained elusive. Here, we address this question by analysing the aggregation kinetics of a large set of self-assembling proteins, from those associated with disease, over those whose aggregates fulfil functional roles in biology, to those that aggregate only under artificial conditions. Remarkably, we find that essentially all systems that assemble by a nucleated-growth mechanism are capable of significant self-replication on experimentally accessible timescales. However, comparing the intrinsic timescales of self-replication with the timescales over which the corresponding aggregates form in a biological context yields a clear distinction; for aggregates which have evolved to fulfil a structural role, the rate of self-replication is too low to be significant on the biologically relevant timescale. By contrast, all analysed proteins that aggregate in the context of disease are able to self-replicate quickly compared to the timescale of the associated disease. Our findings establish the ability to self-replicate as both a ubiquitous property of protein aggregates and one that has the potential to be a key process across aggregation-related disorders.

## Introduction

The self-assembly of proteins into ordered, homo-molecular filaments is a process associated with a range of currently incurable human disorders, from prion diseases, through sickle cell anaemia, to Alzheimer’s disease (*1*,*2*). In such disorders, proteins that are normally monomeric form aggregates, such as the highly stable amyloid fibrils, with often deleterious effects for the organism (*1*). However, filamentous protein self-assembly also plays important roles in a functional context, including in the formation of cytoskeletal structures, such as the polymerisation of actin (*3*). Even amyloid fibrils, which, unlike actin, are generally highly resistant to depolymerisation, are encountered in functional contexts throughout nature, from structural elements in bacterial biofilms, to long-term memory formation in marine organisms (*4*). In addition to these functional and disease associated assemblies, the aggregated state, in the form of amyloid, has been proposed to constitute a general, stable conformation for a large number of proteins (*5*). However, many proteins can only reach this state with the help of perturbations such as shaking, heating or extreme pH conditions due to the large energy barrier that prevents their conversion into amyloid under physiological conditions. With current advances in the theoretical descriptions of aggregation and the increasing accuracy of biophysical measurements in recent years, it is now possible to determine the aggregation mechanisms of many of these proteins (*6*, *7*). Combining the data from dozens of other works, we here determine the mechanism of aggregation for a range of peptides and proteins, in order to elucidate mechanistic commonalities and differences. While various other types of protein aggregates exist, including well-defined macromolecular assemblies, such as virus capsids, or hierarchical assemblies, such as intermediate filaments (*8*), we focus here on those proteins that aggregate into homo-molecular, filamentous aggregates, via a nucleated polymerisation mechanism, primarily amyloids.

## Results

### The presence of a self-replication mechanism leads to fundamentally different aggregation behaviour

In descriptions of linear self-assembly, the underlying processes naturally fall into two categories: *growth processes* which increase the size of existing aggregates, and *nucleation and multiplication processes* which generate new aggregates (*9*). The first category, growth, usually proceeds by addition of monomers from solution to growth-competent aggregate ends, increasing their length, but leaving the total number of aggregates unchanged. The second category, processes that increase the number of aggregates, can be further classified into *primary nucleation* processes, which only involve monomeric protein and are independent of the concentration of aggregates, and *secondary processes* (or *multiplication processes*) which do involve existing aggregates. Primary nucleation can take the form of homogeneous nucleation in solution or heterogeneous nucleation on an interface, whereas secondary processes include the fragmentation of fibrils and the catalysis of nucleation from monomers on the surface of existing fibrils in secondary nucleation. Primary nucleation is always a necessary first step in the formation of aggregates from purely monomeric proteins; by contrast, the presence of secondary processes is not required to fully convert a system to its aggregated state.

The aggregation behaviour of systems with and without secondary processes is fundamentally different (Fig. 1). When secondary processes are active, existing aggregates self-replicate, that is they create new aggregates auto-catalytically, thereby accelerating the overall rate of elongation, which in turn speeds up the production of even more aggregates through secondary processes in an iterative manner. This auto-catalytic feedback loop means that new aggregates amplify very rapidly once they have reached a limiting concentration. The parent-fibril from a single primary nucleation event can thus produce many child-fibrils through self-replication and the aggregate mass increases exponentially with time (see equation (2)). By contrast, in the absence of secondary processes, each aggregate needs to be initiated by a primary nucleation event, which is independent of the amount of fibrils present. The aggregates mass thus increases more gradually and with polynomial scaling in time (see equation (1)). The integrated rate laws describing aggregation under constant monomer conditions are given by

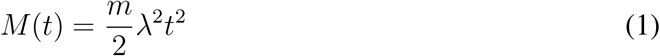

in the absence of self-replication and

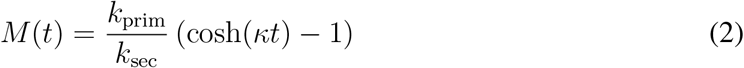

in the presence of self-replication, where *M*(*t*) is the mass concentration of aggregated protein at time *t*, the parameters *κ* and *λ* are defined as 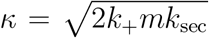 and 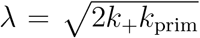, *k*_prim_ and *k*_sec_ are the rates of primary nucleation and secondary processes, respectively, *k*_+_ is the rate constant of growth and *m* is the concentration of monomer (*10*). The rate of secondary processes *k*_sec_ can have contributions from both secondary nucleation and fragmentation and this rate can vary with monomer concentration in different ways, depending on the specific mechanism. More details can be found in Meisl et al. (*7*, *9*).

**Figure 1:**
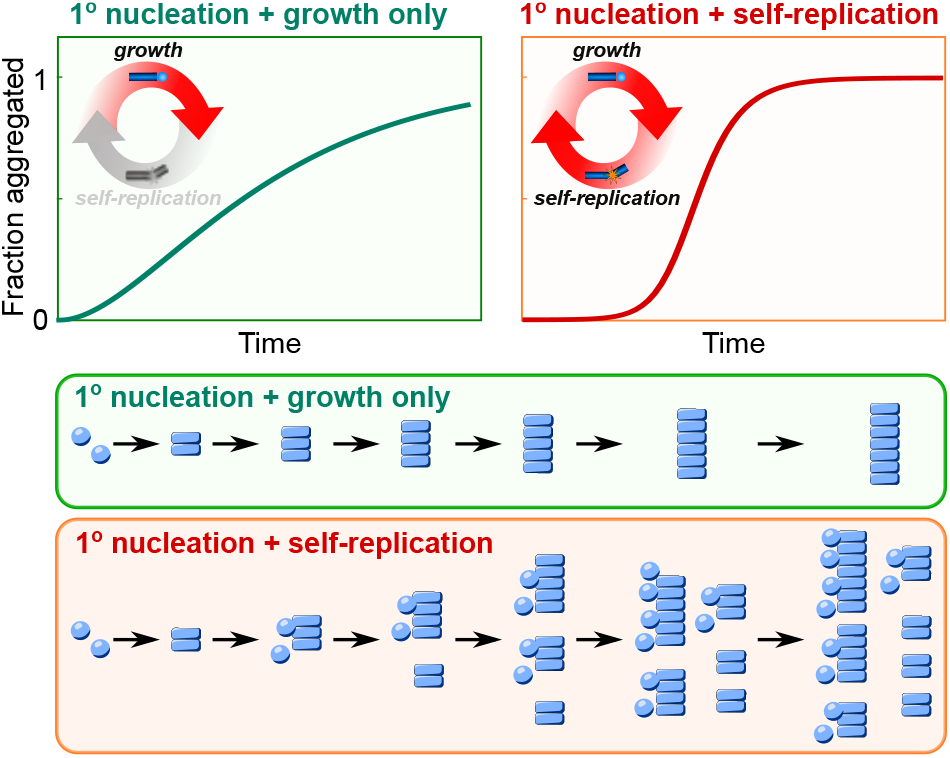
Effect of self-replication. Illustration of the kinetic curves of aggregate concentration over time without (left) and with self-replication (right) along with a schematic of the reaction in both cases. When aggregation proceeds via nucleation and growth only, without self-replication, each primary nucleation event gives rise to only one fibril and the aggregate concentration increases gradually. By contrast, when self-replication is present, here illustrated in the from of secondary nucleation, each primary nucleation event gives rise to many fibrils. The positive feedback loop of self-replication leads to exponential growth of aggregate mass and kinetic curves with a much more sudden increase.

When secondary processes are present, the ability of aggregates to self-replicate in an auto-catalytic manner can make such systems extremely sensitive to the introduction of seed fibrils (Fig. 2g). This property is exploited in several amplification assays, which can amplify a single replication-competent aggregate to macroscopically detectable levels (*11*–*13*). However, in many biological contexts, this susceptibility to amplify small fluctuations of aggregate concentrations may not be a desirable property, making the speed of the formation, as well as the spatial distribution of new aggregates, difficult to control. In the context of disease, the amplification of spontaneously formed or transferred seed aggregates may be crucial in allowing pathology to persist and spread. The effect of small fluctuations in seed concentration on the macroscopic behaviour is demonstrated in Fig. 2d,g for two proteins CsgA and A*β*42. CsgA is an *E. coli* protein that forms functional amyloids during biofilm formation (*14*,*15*) and does not display any significant secondary processes. This is evidenced by the fact that measurements of its aggregation kinetics are well described by a model including only growth and primary nucleation (Fig. 2e) and that the addition of small concentrations of preformed seeds has no effect on its aggregation behaviour (Fig. 2d). A*β*42 is one of the main proteins whose aggregation is associated with Alzheimer’s disease. Its aggregation mechanism contrasts with that of CsgA as it is dominated by the secondary nucleation of monomers on the surface of existing fibrils (*16*). Measurements of its aggregation kinetics cannot be described by a model that does not include a secondary process (Fig. 2h). Moreover, A*β*42 aggregation is very sensitive to the addition of preformed aggregates; even seed concentrations that are orders of magnitude lower than the concentrations of monomeric protein in solution lead to a significant change in the aggregation kinetics (Fig. 2g). With these two proteins exemplifying the two distinct behaviours, we now set out to investigate the universality of self-replication and potential correlations of its presence with the role the protein aggregates play in biology.

**Figure 2:**
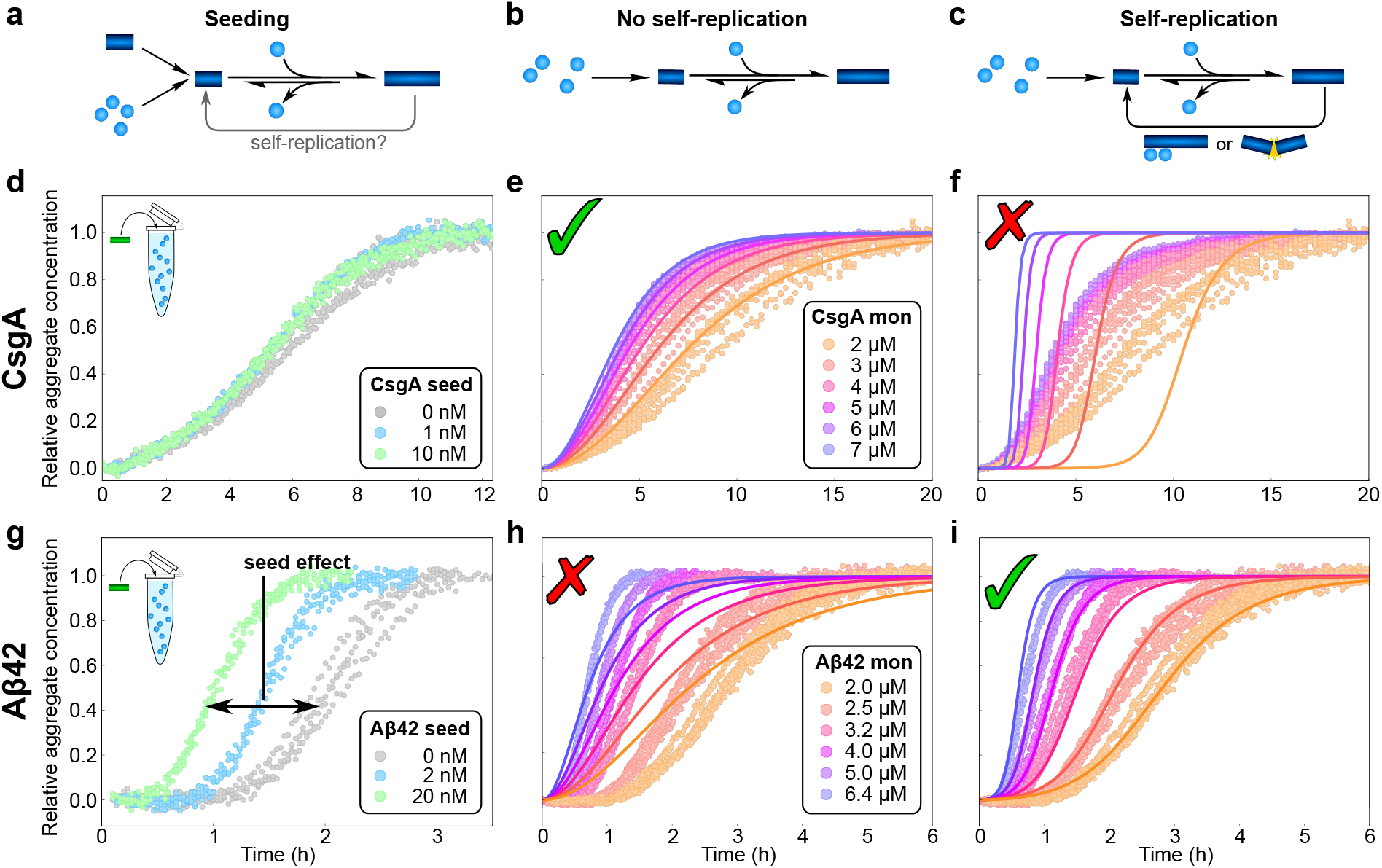
Aggregation kinetics with and without self-replication. **(a-c)** Illustration of the aggregation mechanism used in producing the fits. **(d, g)** Aggregation kinetics of CsgA, 5 *μ*M monomer (d) and A*β*42, 2 *μ*M monomer (g) (*17*), with increasing concentrations of preformed seeds, monitored by Thioflavin T fluorescence, a reporter of the mass of aggregates formed. While the CsgA behaviour is not significantly affected by the presence of seeds, a large effect can be seen for A*β*42 aggregation, even when the seed concentration is up to 3 orders of magnitude below that of the monomeric protein. **(e, f, h, i)** Aggregation kinetics of CsgA (e,f) (*18*) and A*β*42 (h, i) (*19*) at a range of monomer concentrations in the absence of seeds. The solid lines are global fits of the integrated rate laws in the absence (e, h) and presence (f, i) of secondary processes. In (f) a significant contribution of secondary nucleation is enforced to illustrate the misfit. Data are recorded in triplicates at each concentration, all points are shown.

### The ability to self-replicate *in vitro* is a general characteristic of aggregating proteins

Recent advances in chemical kinetics have allowed us to link the macroscopic aggregation kinetics (Fig. 2) to the underlying molecular mechanisms through integrated rate laws for a range of aggregating proteins (*7*, *20*). While the detailed equations depend on the specific aggregation mechanisms, they have several fundamental properties in common. At a given monomer concentration, the aggregation curves are predominantly determined by two parameters: *λ*, a measure for the rate at which primary nucleation contributes new aggregates, and *κ*, a measure for the rate at which secondary processes contribute new aggregates (see also equations (1) and (2)). The relative magnitude of these two rates determines which one of the two processes dominates the overall production of new aggregates. By application of our kinetic analysis framework (*7*), we determined the values of *λ* and *κ* for 7 functional amyloids, 9 pathological amyloids and 8 which do not form amyloid in a biological context (not counting mutants or fragments of the same protein). In order to have the most representative measure of the intrinsic aggregation mechanism of the proteins and avoid bias, we applied stringent selection criteria (see Methods) and, in particular, only used aggregation data under quiescent conditions. Shaking and agitation are commonly used to induce aggregation, but can considerably alter the mechanism (*16*) by promoting fragmentation and inducing aggregation through shearing (*21*). By excluding such datasets we avoid an artificial bias towards fragmentation dominated mechanisms. An example of the analysis performed for all these proteins is shown in Fig. 2, for both the functional amyloid CsgA and the disease-associated A*β*42 peptide. Remarkably, we find that almost all of these systems, regardless of their biological role, display the ability to self-replicate via a secondary process when aggregating *in vitro*. The only exceptions to this are some of the functional aggregates, namely actin, CsgA, and FapC.

The results of our analysis of these data are summarised in Fig. 3 which shows a rate diagram for amyloid forming proteins. The aggregation mechanisms are quantified by the rate at which new aggregates are produced via a primary nucleation pathway, *λ*, and the rate at which they are produced via a pathway involving secondary processes, *κ*. We find that the vast majority of biological protein aggregates, whether functional or disease-associated, are able to self-replicate and are thus located in the upper left hand section of the plot in Fig. 3. Thus, the question arises whether the presence of secondary processes in protein aggregation is the default state, or if only the subset of proteins that is prone to aggregation in biological systems, either in a disease context or as functional assemblies, is biased towards self-replication. To answer this question, we included proteins whose aggregation does not occur in a biological context, but can be triggered by harsh conditions such as low pH and high temperature *in vitro*. These systems also, without exception, display the ability to self-replicate. Thus, we conclude that, if a protein can be made to form filamentous aggregates, the ability to self-replicate appears to be the default state. Only a few functional assemblies appear to have evolved to suppress this property to such a degree that it is no longer readily observable on the timescales of *in vitro* experiments.

**Figure 3:**
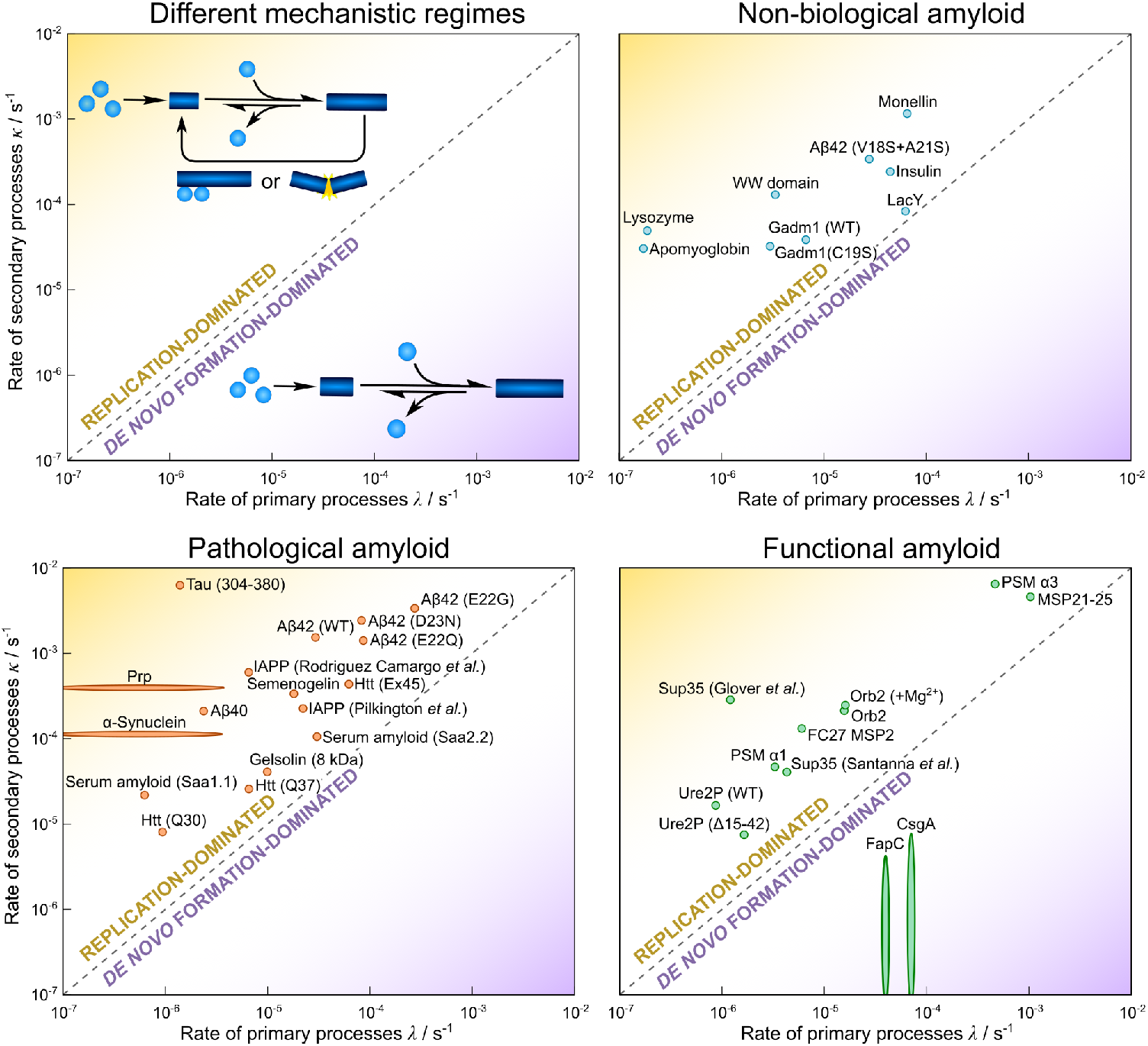
Rate diagrams of aggregation mechanisms show self-replication is ubiquitous. The rate at which new aggregates are produced by secondary pathways, *κ*, is plotted against the rate at which primary pathways produce new aggregates, *λ*. On the dashed line the rates of the two processes are equal. It separates systems dominated by primary nucleation, bottom right corner, from systems dominated by a secondary process, top left corner. In primary nucleation dominated systems, when secondary processes are too slow, only an upper bound for the rate of secondary pathways can be obtained and similarly, when primary nucleation is so slow that seeding is required, only an upper bound for the primary rate can be obtained. These cases are here illustrated by elongated points. Proteins are split into 3 classes: those forming pathological amyloids (bottom left), those forming functional amyloids (bottom right) and those that do not generally form amyloid under physiological conditions (top right). Labels are shown above or to the right of the corresponding data point. Note: A*β*42 (V18S+A21S) is classed as a non-biological amyloid because it is an artificial mutant of A*β*42, itself pathological, designed specifically to try to abolish secondary nucleation, see Thacker *et al*. (*22*). Details on all proteins are given in the SI.

As outlined, there are two distinct molecular mechanisms that can give rise to self-replication *in vitro;* fragmentation of aggregates and secondary nucleation of monomers on the surface of existing aggregates. Fragmentation, as it does not require any specific molecular interactions, may appear an obvious candidate for a default mechanism of self-replication: it tends to be significant under agitation (*16*, *23*, *24*), but has also been found to be important under quiescent conditions in some systems such as yeast prions (*25*). Interestingly, fragmentation induced by the proteasome has also been proposed as an important process of self-replication in cell culture (*26*). However, more surprisingly, secondary nucleation on the surface of existing aggregates has in fact been established as the main mechanism of self-replication under quiescent conditions in many amyloid forming proteins, where this process has been studied in detail (*16*, *27*–*31*). Remarkably, for the A*β*42 peptide, even attempts to specifically abolish the ability to secondary nucleate via targeted mutations were unsuccessful. The mutants displayed significant changes in fibril morphology, but self-replication still proceeded by secondary nucleation (*22*) (see also Fig. 3: A*β*42 V18S+A21S). Secondary nucleation is not exclusive to protein aggregation and appears in many other contexts, such as crystal growth, where it is often a result of strain and local defects (*32*–*34*). While our data are too limited to clearly establish the specific mechanism of self-replication for most amyloid-forming systems, the ubiquitous presence of self-replication, alongside the fact that assembly of most of these structures induces strained conformations (*35*), make for an intriguing correlation. In addition to fragmentation, secondary nucleation in particular, not just self-replication in general, may thus be a key process in the formation of many filamentous aggregates.

We further note that the specific values of the rates *λ* and *κ* fall into a relatively narrow range here, which likely reflects the experimental limitations of measuring aggregation kinetics: in order to be observable with standard techniques on an experimentally feasible timescale, the conditions and concentrations will be adjusted to result in aggregation over the course of minutes to hours. However, while the absolute rates are biased by the experimental limitations, the dominance of secondary over primary processes remains a robust finding.

### The timescales of self-replication correlate with biological roles

To further investigate the importance of our findings in the context of the respective biological systems in which these proteins are found to aggregate, we investigated the potential significance of self-replication at relevant biological timescales and concentrations. While secondary processes, such as fragmentation and fibril-catalysed nucleation, conceivably proceed in a similar manner in biological systems as they do *in vitro*, the process of primary nucleation is likely to differ more significantly. *In vitro*, air-water interfaces (*36*) or the surface of the reaction vessel may serve as heterogeneous primary nucleation sites, whereas *in vivo* nucleator proteins for functional aggregates (*37*), or lipid membranes for disease-associated ones (*38*) may trigger primary nucleation. To assess the potential role of self-replication in a biological context, we therefore focus exclusively on the rate constant of self-replication obtained from kinetic analysis. The aim of this comparison is to answer the question whether the intrinsic self-replication propensity, as measured *in vitro*, correlates in any way with the roles that protein aggregation plays in living systems. Generally, aggregation is slowed in living systems, for example through the action of chaperones or of active removal processes (*39*,*40*). Thus, our analysis investigates if, despite these *in vivo* effects, the intrinsic self-replication propensity can be a predictor of disease association.

For the subset of functional and disease-associated proteins analysed in Fig. 3 for which a set of aggregation data at varying monomer concentrations is available, we determined the rate constants and reaction orders. Thus, we were able to evaluate the time to double the number of aggregates through self-replication, *t*_2_ = ln(2)/*κ*, at the protein concentrations encountered in the respective *in vivo* environment. We then compared this timescale to the characteristic timescale of the *in vivo* process in which aggregation takes place, such as the time for biofilm maturation for CsgA or the disease duration for the prion protein, see Fig. 4.

**Figure 4:**
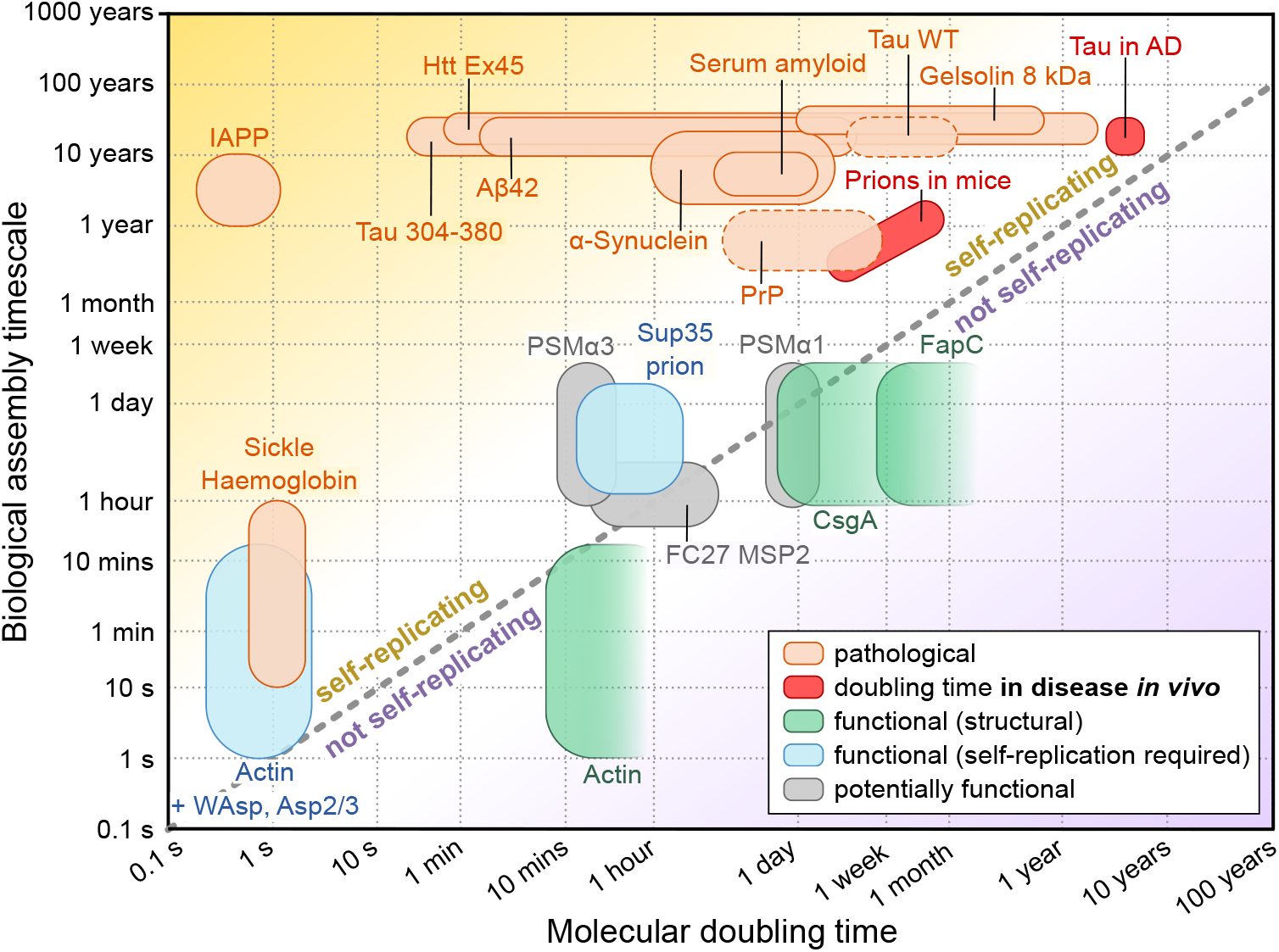
Timescales of self-replication and relevant in vivo timescale. In the bottom right half, the timescale of self-replication is longer than the relevant *in vivo* timescale, making self-replication unlikely to be able to contribute to the aggregation kinetics *in vivo*. By contrast, in the top left half, the timescale of self-replication is much shorter than the relevant *in vivo* timescale, making a contribution of self-replication to the process likely. The timescales of self-replication were computed using the fitted rate constants and the protein concentrations in their respective *in vivo* environments, the latter being the main source of uncertainty. For two systems, prions in mice and tau in Alzheimer’s disease (AD), the self-replication rates have been determined in the *in vivo* system directly, requiring no further computation or knowledge of the *in vivo* concentrations (red). For FapC, CsgA and actin, the experimental aggregation kinetics are described well with a mechanism that does not include self-replication, therefore we can only obtain a lower limit on the timescale for those proteins. Dashed lines for PrP and tau denote mild shaking conditions, for details see Methods.

Remarkably, we find that all disease-associated proteins, without exception, show self-replication timescales which are much shorter than the timescales of the associated disease. Therefore, all disease-associated aggregates posses the intrinsic ability to replicate sufficiently quickly for self-replication to be a relevant mechanism in disease progression. Rates of self-replication measured in the relevant system *in vivo* are very rare, but we were recently able to determine them in two systems, prions in mice (*41*) and tau in Alzheimer’s disease (*42*). These rates are included here alongside the rates calculated from *in vitro* measurements to serve both as a comparison to the *in vitro* numbers and as validation of our conclusions on the potential importance of self-replication. While replication is somewhat slowed compared to the *in vitro* behaviour of the same protein, both systems are still clearly in a regime dominated by the self-replication process.

By contrast, the functional assemblies cluster much closer to the threshold at which self-replication becomes too slow to be relevant on the biological assembly timescale (diagonal dashed line in Fig. 4). For those functional assemblies that have been established to fulfil structural roles, FapC, CsgA and actin, the archetypal biological structural element, the self-replication timescale exceeds that of the relevant *in vivo* process, suggesting that self-replication is too slow to be relevant *in vivo*. The timescales of self-replication appear to be just long enough to have no significant contribution to the aggregation process on the relevant biological timescale. This reduction of self-replication only by the minimal amount necessary, is further indication in support of the idea that self-replication is ubiquituous and has to be selected against by evolution if undesirable. Remarkably, the two functional systems for which self-replication is likely to be a desirable property, yeast prions (Sup35) and actin in presence of a branching agent WAsp, Arp2/3, are indeed situated in the region of the plot where self-replication is significant. Specifically, the biological role of yeast prions is believed to involve the transfer of the prion between yeast cells, thus requiring self-replication of the aggregates on timescales relevant for yeast reproduction. In actin assembly, if branching, a secondary process, is required, it can be induced in actin in a controlled manner by other molecules such as Wiskott-Aldrich syndrome protein, WAsp (*43*) (see Fig.4). Finally, the PSMs, expressed by Staphylococcus bacteria, and FC27 MSP2, expressed by the malaria parasite, form amyloid fibrils under native conditions *in vitro*, but the specific biological roles of these assemblies, and thus whether one would expect their self-replication to be a desirable property, have yet to be established.

Thus, while there appears to be some evidence for an evolutionary pressure to prevent self-replication if aggregates are involved in certain functions, it is unclear how this is achieved. Intriguingly, modification of some structural functional amyloids, e.g. by removal of regions from the sequence, can lead to increased rates of self-replication: CsgA and FapC consist of multiple copies of imperfect repeats of 20-35 residues (*14*, *44*). Each repeat is predicted to fold into a beta-hairpin conformation which forms a self-contained rung in a beta-helix structure (*45*,*46*) and thus constitutes a readily available building block for the efficient construction of an amyloid edifice. Stepwise removal of these repeats increases the tendency of the fibrils to fragment (*47*). Thus, accumulation of repeats within the protein sequence, at least in these bacterial amyloids, appears to suppress self-replication over primary nucleation. Beyond this observation, a further investigation of the proteins analysed here for patterns in their sequence did not reveal any clear features associated with the propensity to self-replicate (see SI Fig. S23) (*48*). The ubiquity of self-replication may prevent a specific sequence feature from being identified across the diverse set of proteins analysed here. As the set of proteins with known aggregation mechanisms increases, further stratification of the data to investigate the presence of features important for self-replication in different classes of proteins may become possible.

## Discussion

In conclusion, our results provide insights into both the widespread presence of self-replication in amyloid formation, and the potential importance of this process in disease. Almost all aggregating proteins studied have the ability to self-replicate, although the propensity to do so appears to have been deselected for by evolution in some, but not all, proteins that have evolved to aggregate in a functional context. In light of these results, we propose that for certain functional roles, such as structural support, a crucial aspect in determining whether a protein is suitable for the formation of functional structures may be its propensity to self-replicate. In addition to making the system susceptible to fluctuations in initial seed concentration, secondary nucleation would lead to the formation of new fibrils along the length of existing fibrils, making the control over the location of these new fibrils difficult. Similarly, a high propensity to fragment is unlikely to be desirable in a structural context.

By contrast, the ability of aggregates to self-replicate appears to be a central prerequisite for their involvement in disease. Indeed, prion disease, the archetypal protein aggregation disease, requires the self-replication of its disease-causing aggregates to propagate between individuals (*49*). Moreover, in many model organisms of neurodegenerative disease it has been shown that the introduction of seeds can trigger the formation of new aggregates (*50*, *51*), leading to the description of several such diseases as prion-like (*52*). In recent work, we established the replication mechanism of prions in mice and found that it is in fact consistent with the self-replication mechanism of PrP aggregates *in vitro* (*41*). In another study, we showed that the relative difference in the replication rate of two strains of *α*-synuclein aggregates produced *in vitro* was mirrored in the survival times of mice infected with these strains (*53*). Finally, in recent work we established that self-replication is the rate-limiting step of the accumulation of tau aggregates in Alzheimer’s disease (*42*). In light of these clear connections between the *in vitro* mechanisms and the mechanisms active in disease, our finding of a remarkable correlation between replication timescales and disease association indicates that self-replication, by the simple mechanisms that act on purified proteins *in vitro*, should be considered a key factor in the pathology of a wide range of aggregation-related disorders.

## Material and Methods

### Choice of data for kinetic analysis

A large number of publications exist that contain largely qualitative data of the aggregation of proteins. To also be suitable for interpretation in the context of a quantitative kinetic analysis as we present it here, a number of conditions need to be met. In particular we **excluded**

- experiments which did not measure a reporter of the mass of protein aggregates formed,
- datasets which showed aggregation that was so fast that nucleation takes place fully in the dead-time of the experiment (evident by a lack of positive curvature of the kinetic curves),
- datasets in which for any other reason the purity of the sample was questionable,
- kinetic experiments that were performed under significant agitation (as that biases the system towards fragmentation (*16*)). Brief shaking prior to measurement, e.g. 5 seconds every 10 minutes as for gelsolin (*54*), was deemed acceptable.

The two exceptions to the last point are the data for full length tau and PrP, both of which were obtained under mild shaking. The presence of shaking may lead to a slight underestimation of the timescale of self-replication. However, we are confident for both of these systems self-replication is still sufficiently rapid to be relevant *in vivo*, because we have in fact obtained measures for the timescale of replication directly from data in living systems.

The biggest experimental problem is the presence of small amounts of preformed aggregates at the start of the experiment. While impurities other than the protein of interest are usually no longer an issue after being removed by standard purification protocols, seed aggregates can easily reform after purification due to improper handling during sample preparation that induces nucleation. The presence of such seeds can lead to a significant shortening of the lag phase in systems dominated by secondary processes. As this might lead to misinterpretation of these systems as lacking secondary processes, this was a major concern in our analysis and the analysed datasets were carefully inspected to minimise the risk of such a situation. In the light of the overwhelming evidence for secondary processes, together with the fact that seed impurities would bias the analysis towards a more primary nucleation dominated conclusion, we conclude that this source of error is unlikely to be significant in the datasets selected here.

### Protein purification, assay conditions and biological role

Please refer to the original publications for the detailed methods of protein purification and conditions of the aggregation assay. In the SI, we give the original source for every dataset used, an overview of the biological roles of each of the studied proteins and the rationale behind the choice of relevant biological timescale.

### Fitting of kinetic data

The aggregation kinetics were fitted using the AmyloFit interface at www.amylofit.com (*7*). All data were fitted with a model which includes the processes of primary nucleation, elongation and secondary nucleation, given by the equations

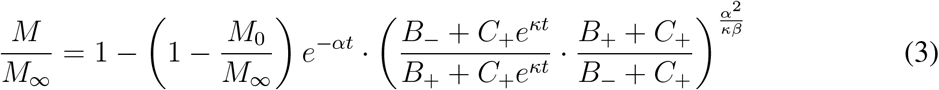

where the definitions of the parameters are

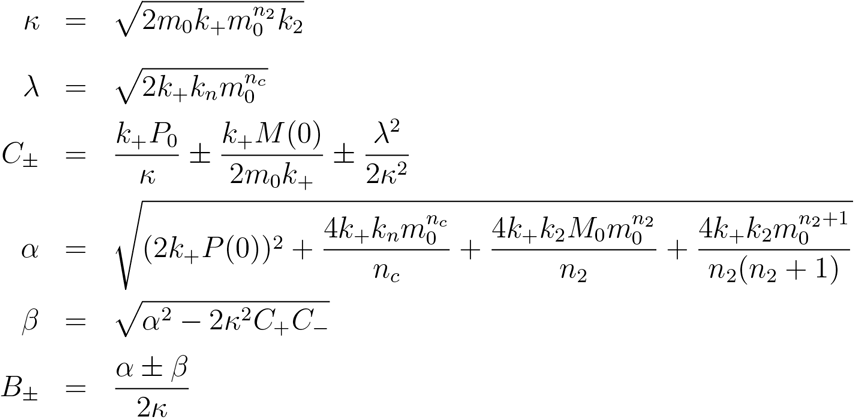

where *M*(*t*) is the mass concentration of aggregates, *m*_0_ is the monomer concentration at the beginning of the aggregation reaction, *k_n_*, *k*_+_ and *k*_2_ are the rate constants of primary nucleation, elongation and secondary nucleation, respectively, and *n_c_* and *n_2_* are the reaction orders of primary and secondary nucleation, respectively. In some special cases, the two step nature of secondary nucleation (for some datasets of A*β* and tau) or of elongation (for *α*-synuclein) had to be taken into account explicitly, see Meisl *et al.* (*7*, *9*) for details on the refinement of the model in those cases. The details for all proteins can be found in the SI.

When data at a range of monomer concentrations are available, the models can be fitted to extract both the reaction orders as well as the rate constants. These parameters can then be used to calculate the rates *λ* and *κ* also at monomer concentrations not directly measured in the experiment. When data at only one monomer concentration is available, only *λ* and *κ* can be determined, but the values of the reaction orders and reaction rate constants cannot be determined separately. Thus, only those proteins for which for which data at a range of monomer concentrations has been measured can be used to calculate relevant timescales at *in vivo* concentrations, reflected in the fact that Fig. 4 contains only a subset of the proteins shown in Fig. 3. For each system we determined either the rate constants and reaction orders, *k*+*k_n_*, *k*_+_*k*_2_, *n_c_* and *n*_2_ (if data at a range of monomer concentrations were available) or only the rates *λ* and *κ* (if there were insufficient data to determine the reaction orders and rate constants separately). When data at multiple concentrations was available, we chose an intermediate concentration to calculate *λ* and *κ* for Fig. 3.

For systems not displaying any detectable secondary processes (e.g. CsgA), we set a conservative upper bound for the rate of self-replication as 10% that of primary nucleation, i.e. *κ* ≤ 10*λ*. In those cases where no self-replication was detectable in experiments, it was also not possible to determine the reaction order of the secondary process. Therefore, in order to calculate a bound for the self-replication rate at *in vivo* concentrations, we used *n*_2_ = 0. This choice was made to ensure that the quoted rate is still an upper bound on the true rate of self-replication; as the monomer concentrations in the *in vitro* experiments are higher than those encountered *in vivo*, extrapolation of the rates determined in experiment to *in vivo* concentrations will yield the maximal rate of self-replication for the minimal choice of reaction order, *n*_2_. As this only concerns systems that are in the lower right region of Fig. 4, an upper bound on the self-replication rate (corresponding to a lower bound on the doubling time), is all that is needed.

A similar problem is encountered for systems where primary nucleation is so slow that to observe aggregation on experimentally accessible timescales seeded experiments have to be used (e.g. *α*-synuclein). In those cases we obtain an upper bound for the rate of primary nucleation by calculating the initial rate of nucleation from secondary processes acting on the seeds and set the primary rate to be equal to this (i.e. the upper bound for primary nucleation is that it produces as many nuclei as secondary processes at the beginning of the seeded reaction). Expressing this in terms of the rate constants gives 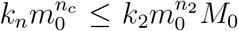, which can be used to show that 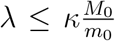, where *m*_0_ and *M*_0_ denote the initial monomer and fibril concentrations, respectively. As primary nucleation is not considered in Fig. 4, an estimate of the reaction order is not required.

### Calculation of timescales

The biologically relevant timescale for diseases was chosen to be the approximate time from diagnosis/symptom onset to death, with the exception of sickle cell anaemia, for which the relevant timescale was chosen to be the speed of onset of a sickle cell crisis. For the functional proteins, biologically relevant timescale is the timescale over which the associated process takes place i.e. the timescale of biofilm assembly (CsgA, FapC), of colony spreading (PSM*α*), of cytoskeleton assembly (actin), of yeast cell multiplication (Sup35 prions), and of formation time of the merozoite form of the malaria parasite (FC27 MSP2). The molecular doubling time was computed using the rate constants and reaction orders determined in our fits of *in vitro* data along with estimates of the *in vivo* relevant concentrations of the aggregating proteins. As kinetics at a range of different protein concentrations was available only for a subset of the data analysed, the timescales at *in vivo* relevant concentrations could be determined only for this subset of the data, thus not all proteins shown in Fig.3 could be included in Fig. 4. Detailed numbers and references are given in SI tables 1-3.

## Supplementary materials

Materials and Methods

Supplementary Text

Figs. S1 to S23

Tables S1 to S3

## Acknowledgments

We thank our colleague and friend the late Prof Sir Christopher Dobson for his insightful comments and helpful discussions. We also thank Prof Margaret Sunde, Prof Robin Anders and Dr Christopher Adda for their expert input.

## Funding

This work was supported by the Danish Council for Independent Research—Natural Sciences (FNU-11-113326) (MA), Peterhouse College, Cambridge (TCTM), Sidney Sussex College, Cambridge (GM), the European Research Council Grant Number 669237 (DK, GM), and a Herchel Smith Research Studentship (CKX).

## Author contributions

GM and CKX performed literature data extraction, analysis and visualisation. JDT performed the CsgA experiments. GM wrote the manuscript. All authors edited the manuscript.

## Competing interests

GM is a scientific consultant for Wren Therapeutics.

## Data and materials availability

All data needed to evaluate the conclusions in the paper are present in the paper and/or the Supplementary Materials.

## Notes

### Competing Interest Statement

The authors have declared no competing interest.

