## Supplementary Information for "Uncovering the universality of self-replication in protein aggregation and its link to disease"

### 1 Aggregation mechanisms

In the past, we have developed a range of integrated rate laws for variations on the mechanism of aggregation, for example due to differing reaction orders and saturation of particular processes. Details on these mechanisms can be found in Meisl et al. [1]. The majority of proteins in this work, including all for which only one concentration was available, were analysed with a model including primary nucleation, single step secondary nucleation and elongation. The proteins analysed with a different model are A$\beta$ and $\alpha$-synuclein.

### 2 Proteins

Here we briefly discuss each protein studied, including the biological role of aggregation and where the data were obtained. For most proteins, the data have not been previously analysed by fitting of integrated rate laws to obtain reaction rates. In those cases, i.e. when data was analysed for the first time in this work, the fits are shown in Figs. S1 to S22. For datasets which had been previously analysed (A$\beta$ and some tau and IAPP datasets) please refer to the cited works for the details of the fits.

#### 2.1 α-Synuclein

α-Synuclein in its amyloid fibril form is the major protein component of the Lewy bodies and neurites that are one of the hallmarks of Parkinson's disease [2]. Gaspar et al. investigated the aggregation of full-length wild type α-synuclein [3]. Data analysed in our study are those shown in figure 2c of that paper. The biologically relevant timescale is the length of Parkinson's disease.

## 2.2 Aβ

Aggregation of both isoforms of the Aβ peptide, $A\beta_{40}$ and $A\beta_{42}$, is associated with Alzheimer's disease (AD) [4, 5]. The point mutations E22G, E22Q, and D23N are all associated with familial AD. The rate constants for $A\beta_{40}$ are from Meisl et al.[5]. The $A\beta_{42}$ values, including all familial AD-associated mutants and the data and fits shown in Fig. 1 of the main text are from Yang et al. [6] (data and fits shown in figure 5 of that paper). The seeded data shown in Fig. 1 are from Weiffert et al. [7]. The V18S+A21S $A\beta_{42}$ is an artificial variant which was designed to try to abolish secondary nucleation, rate constants for this are determined in Thacker et al. [8]. For A$\beta$42, while the aggregation at $\mu$M concentrations, under physiological slat concentrations, is dominated by secondary nucleation with a reaction order of 2, there appears to also be fragmentation of fibrils that proceeds at a much lower rate than secondary nucleation, as shown in Meisl et al. [9]. However, when $\kappa$ is calculated at low concentrations, one has to take into account the contribution from fragmentation as this process is expected to dominate over seocndary nucleation at sufficiently low concentrations, due to the fact that the fragmentation rate does not scale with monomer, unlike the secondary nucleation rate. We thus also evaluate $\kappa$ due to fragmentation, and use this to construct the plots in Fig. 3. The relevant rate constants are given in table 3 under "Aβ42 (fragmentation)". The biologically relevant timescale is the length of Alzheimer's disease (specifically average time from Braak stage 3 to Braak stage 6).

#### 2.3 Actin

Actin is one of the major components of the eukaryotic cytoskeleton, and its conversion between monomeric (G-actin) and filamentous (F-actin) states facilitates the regulation of a multitude of cellular functions and properties, including cell shape and motility [10]. Actin polymerisation is a distinct process from amyloid aggregation, with F-actin lacking the canonical cross-β structure [11, 12]. Pure actin is only able to form polymers, and requires actin binding proteins (ABPs) to nucleate new F-actin fibres from existing F-actin surfaces, which we can equate to amyloid self-replication processes [13]. Higgs et al. investigated the kinetics of actin polymerisation in the absence and presence of ABPs. The ABPs under study were the Arp2/3 complex, a weak cellular nucleating factor, and WA, which stimulates the Arp2/3 nucleation ability and consists of the C-terminal regions of Wiskott-Aldrich syndrome protein, N-WASP, and three isoforms of Scar, collectively termed the WASp/Scar protein family [14]. Data analysed in our study were extracted from figure 9 [14]. In the tables, aggregation under these conditions is referred to as "Actin + WAsp". The biologically relevant timescale is the length of cytoskeleton formation / rearrangement.

#### 2.4 Apomyoglobin

The aggregation of apomyoglobin is not related to any human diseases, nor does it seem to play a role *in vivo*. The ability of apomyoglobin to form amyloid fibrils is therefore used as a model system and proof of the apparent universality of amyloid aggregation. Tryptophan residues at positions 7 and 14 were found to play an important role in the folding of apomyoglobin [15], and the double apomyoglobin mutant W7F/W14F (W7FW14F) was furthermore found to increase the aggregation propensity of the protein [16]. Unlike the WT protein, the W7FW14F readily aggregates at physiological pH and temperature; Vilasi et al. therefore used this construct in their study[17]. Data analysed in our study were extracted from figure 3 [17].

#### 2.5 CsgA

CsgA is the protein subunit which forms the curli amyloid fibrils in *Enterobacteriaceae* [18, 19]. The curli fibrils are the major protein component of the extracellular matrix and thus play a key role in biofilm structuring. Andreasen et al. investigated the aggregation of the recombinantly expressed mature form (residues 22-151) of CsgA [20]. The data shown here, including the data and fits shown in Fig.1 of the main text, are from Andreasen et al. [20] (data and fits shown in figure 3a of that paper). The seeded data shown in Fig. 1 have not been published previously. The biologically relevant timescale is the timescale of biofilm formation.

#### 2.6 FapC

FapC is produced by *Pseudomonas* strains and, in its amyloid form, contributes to biofilm structuring and strengthening [21]. The data shown here are from Andreasen et al. [20] (data and fits shown in figure 3b of that paper). The biologically relevant timescale is the timescale of biofilm formation.

#### 2.7 Gadm1 (β-Parvalbumin)

β-Parvalbumin fibrils do not play a known functional or pathological role [22]. Amyloid fibrils of β-parvalbumin are of interest because there is evidence suggesting that they are responsible for food allergies [22]. The aggregation of Atlantic cod β-parvalbumin, Gadm1, was studied by Castellanos et al. [23]. In this study, the aggregation behaviour of wild-type and two variants (I12C and C19S) of Gadm1 were investigated. These mutations were chosen, since residues I12 and C19 may play a role in amyloid assembly and calcium ion binding properties [23]. Data analysed in our study were extracted from figure 3A [23].

#### 2.8 Gelsolin

The aggregation of mutant gelsolin, with the D187N/Y mutation being the most common, is associated with gelsolin amyloid disease, or familial amyloidosis of Finnish type (FAF). Gelsolin is cleaved into several fragments, including 5 kDa and 8 kDa fragments, both containing the D187N/Y mutation. Both fragments are amyloidogenic, with the 8 kDa fragment comprising the majority of FAF-associated fibrils. Solomon et al. characterised the aggregation of these two fragments with the D187N mutation [24]. Data analysed in our study were extracted from figure 1D of that paper, where aggregation was performed at pH 7.4. The biologically relevant timescale is the length of Finnish gelsolin amyloidosis.

#### 2.9 Huntingtin and polyQ peptides

Ectended CAG repeat sequences in trinucleotide (CAG) repeat expansion disorders result in the production of proteins with extended polyglutamine (polyQ) tracts, whose aggregation is characteristic of such diseases [25]. Kakkar et al. studied the aggregation of a peptide containing a polyQ tract of length 45 (sequence $GAMKSFQ^{45}F$) and 14 kDa Huntingtin exon 1 construct, again with a 45Q tract [25]. Data analysed in our study were extracted from figures 1A and S1B-C [25]. Kar et al. explored the effects of polyQ tract length on aggregation kinetics; we analysed their data on the N-terminal region of the protein ataxin-7 with a 30Q tract, and on $SFQ_{37}P_{10}K_2$, a fragment of the Huntingtin N-terminal peptide [26]. Data analysed in our study were extracted from figures 1C and 3B [26]. The biologically relevant timescale is the length of Huntington's disease.

#### 2.10 Islet amyloid polypetide precursor

The aggregation of islet amyloid polypeptide precursor (IAPP) is associated with Type II diabetes, in which aggregates of IAPP are found in the extracellular space of the islets of Langerhans [27]. Pilkington et al. investigated the aggregation of the 37-residue peptide KCNTATCATQRLANFLVHSSNNFGAILSSTNVGSNTY with a disulphide bridge between residues 2 and 7 [27]. Data analysed in our study were extracted from figure 1A [27]. Additionally, both rates and reaction orders for IAPP aggregation were obtained by Rodriguez-Camargo et al. [28], also included here. The biologically relevant timescale is the timescale of Type 2 diabetes onset.

#### 2.11 Insulin

The aggregation of insulin serves neither a functional purpose nor is it associated with pathology. Nielsen et al. studied the aggregation of bovine insulin induced by low pH [29]. Data analysed in our study were extracted from figures 2A-B and 6 [29].

#### 2.12 LacY

The aggregation of the membrane protein LacY serves neither a functional purpose nor is it associated with pathology. Stroobants et al. monitored the aggregation of the wild type LacY protein [30]. Data analysed in our study were extracted from figure 3A [30].

#### 2.13 Lysozyme

The aggregation of hen egg white lysozyme (hewL) serves neither a functional purpose nor is it associated with pathology, instead being used as a model system by which to study aggregation. Hasecke et al. monitored the aggregation of wild-type hewL [31]. Data analysed in our study were extracted from figure 2C [31].

#### 2.14 Monellin

The aggregation of the plant protein monellin serves neither a functional purpose nor is it associated with pathology [32]. Konno et al. studied the aggregation of the wild-type protein [32]. Data analysed in our study were extracted from figure 5A [32].

#### 2.15 MSP2

Merozoite surface protein 2 (MSP2) is a 30 kDa protein found on the surface of the malaria-causing *Plasmodium falciparum* [33, 34]. The structure of MSP2 in this physiological has not yet been fully established, however, some evidence suggests the presence of amyloid-like oligomers, and the ability of MSP2 to form fibrils may play a role in immune recognition [33, 35]. Adda et al. studied the aggregation kinetics of MSP2 from the FC27 strain with a C-terminal hexa-His tag [35]. Data analysed in our study were extracted from figure 5 [35]. Low et al. characterised the aggregation properties of the N-terminal region (MSP1-25), which is conserved between the two main strains, 3D7 and FC27. In addition to the WT fragment, they also studied the aggregation of the [Y7A, Y16A] double mutant, to probe the effects pf tyrosine residues on aggregation. Data analysed in our study were extracted from figure 6 [33]. The biologically relevant timescale is the timescale of merozoite formation. The bounds on biologically relevant concentration was estimated from merozoite numbers and sizes formed in the parasituous vacuole.

#### 2.16 Orb2

Aggregation of Orb2 into amyloid fibrils is important for its function in long-term memory formation in *Drosophila melanogaster*, and its analogues play similar roles in *Aplysia californica* and mice [36, 37, 38]. In *Drosophila melanogaster*, Orb2 is present in two isoforms, the 551-residue Orb2A and the predominant 704-residue Orb2B, which differ only at their N-termini. Although less abundant, Orb2A is required to initiate fibril formation [38, 39]. Bajakian et al. investigated the aggregation of the first 87 residues of Orb2A (Orb2A87), whose metal binding properties may play a role in regulating aggregation [39]. Data analysed in our study were extracted from figure 6A [39].

#### 2.17 Phenol soluble modulins

Phenol soluble modulins (PSMs) are a family of peptides produced by *Staphylococcus aureus* with a range of functions, behaving as toxins, pro-inflammatory agents, or structural entities in biofilms [40]. There are two classes of PSMs, $\alpha$and $\beta$[41]. PSMs of type $\beta$ have been found to be consistently less cytotoxic than the $\alpha$-type peptides, and are likely to play a key role in biofilm structuring [42, 41, 43, 44]. PSM$\alpha$1 fibrils have also been found to promote biofilm stability, whereas fibrillation of PSM$\alpha$3 increases its toxicity, suggesting that its main role may be as a toxin against human cells [45, 46, 47]. Data analysed in our study are those shown in Fig. 1 of Zaman et al.[48]. The biologically relevant timescale is the timescale of biofilm formation.

#### 2.18 Prion protein

Aggregation of prion protein (Prp) is responsible for prion disease and transmission. In vitro rate constants used here are from Sang et al.[49]. The biologically relevant timescale is the length of prion disease. Prp is one of two systems for which we have determined the self-replication rate also in vivo [50]: this was measured in 4 mouse lines which express different concentrations of Prp (resulting in different self-replication rates) and also have different survival times. This variation of self-replication rate, along with the relevant timescale, reflected in the diagonal region marked for ”Prions in mice” in Fig. 3 of the main text.

#### 2.19 Semenogelin

The deposition of aggregated semenogelin-1 (Sg1) protein in seminal vesicles is associated with senile seminal vesicle amyloidosis (SSVA) [51, 52, 53]. Frohm et al. studied the aggregation of a peptide derived from Sg1 with the sequence GSFSIQYTYHV, corresponding to resides 38-48 in in the 439-residue Sg1 [54, 55]. Data analysed in our study were extracted from figure 5 [55].

#### 2.20 Serum amyloid

SAA (serum amyloid A) protein is overexpressed during inflammation and constitutes the major component of the amyloid deposits in amyloid A (AA) amyloidosis [56, 57]. SAA exists in several isoforms, with SAA1, specifically the SAA1.1 isoform, being the most abundant in aggregates in mouse models [58]. Srinivasan et al. investigated the aggregation of the murine SAA1.1 isoform [59]. Data analysed in our study were extracted from figure 3A [59]. The murine SAA2.2 isoform shares 94% sequence identity with the pathogenic SAA1.1 isoform, yet is not pathogenic, despite readily forming fibrils *in vitro* [60, 61]. Ye et al. characterised the aggregation of the murine SAA2.2 isoform [62]. Data analysed in our study were extracted from figure 2 [62]. The biologically relevant timescale is the length of amyloid a amyloidosis.

#### 2.21 Sickle haemoglobin

Under low oxygen conditions, sickle haemoglobin (HbS) polymerises [63, 64]. While not considered to be amyloid aggregation, the mechanistic steps involved in HbS polymerisation Data analysed in our study were extracted from Ferrone et al. [65] figure 9. Sickle haemoglobin has been established to have a very high reaction order; a full model is complex and discussed in Ferrone et al.[66]. For consistency we fit the same model as for the other proteins in this work, and use only the lowest experimentally measured concentration (3.95 $\mu$M). This concentration is also a good lower bound for the in vivo relevant concentration, thus not requiring knowledge of the exact reaction order (and how it may vary with concentration) to calculate $\kappa$ at a different concentration. We therefore obtain directly a lower bound for $\kappa$, i.e. an upper bound for the molecular doubling time, from the analysis of one in vitro concentration, which is sufficient to unambiguously locate sickle haemoglobin in the upper left half of Fig. 3 of the main text. The biologically relevant timescale is the average blood circulation time and speed of onset of a sickle cell crisis.

#### 2.22 Sup35

The *Saccharomyces cerevisiae* Sup35 protein (Sup35p), in its soluble form, is a translation termination factor [67]. Sup35p also exists in a transmissable amyloid form, wherein it functions as a yeast prion, being responsible for the [*PSI*$^+$] phenotype, increasing the phenotypic diversity and thus conferring a survival advantage in changeable environments [68, 69]. The [*PSI*$^+$] phenotype can arise spontaneously and can be propagated to daughter cells via the cytoplasm, dependent on the presence of aggregated Sup35p seeds; self-replication of seeds is therefore crucial for its function [69]. Sant'anna et al. characterised the aggregation kinetics of the 21-residue prion-forming domain of Sup35p (RGNYKNFNYNNNLQGYQAGFQ), corresponding to residues 98-118 of the 685-residue protein [70]. Data analysed in our study were extracted from figure 6 [70]. Glover et al. studied the aggregation properties of the N-terminal and middle (NM) region of the protein, encompassing residues 1-253; this fragment is sufficient to induce the [*PSI*$^+$] in yeast [71]. Data analysed in our study were extracted from figure 4A-B [71]. The relevant timescale is the average time per generation in growing yeast cells.

#### 2.23 Tau

The aggregation of tau is associated with a range of disorders, of which Alzheimer's disease is the most prevalent. The in vitro rates of tau(304-380) are from Rodriguez-Camargo et al. [72], the tau WT rates are from Kundel et al. [73]. The tau datasets from Kundel et al.[73] use heparin to artificially induce primary nucleation, hence the primary rate constants and reaction orders observed in that experiment are not used here. As this means lambda is difficult to estimate, we show these tau data only in Fig. 3 (which is independent of lambda) and not Fig. 2 (this only contains the other tau dataset from Rodriguez-Camargo et al. [72]). While we can't estimate lambda for full length tau, the fact that no aggregation of full length tau is seen in the absence of an inducer of primary nucleation confirms that indeed secondary processes dominate over primary ones also in the full length system. Tau is one of two systems for which we have determined the self-replication rate also in vivo, see Meisl et al. [74]: this was measured in humans with Alzheimer's disease. The relevant timescale is the length of Alzheimer's disease (specifically average time from Braak stage 3 to Braak stage 6).

##### 2.24 Ure2p

The Ure2 protein (Ure2p) is a yeast prion, which behaves similarly to Sup35p [75, 69]. Ure2p is responsible for the [*URE3*] phenotype in its prion form. In its soluble form, Ure2 is a regulator of nitrogen catabolism, which prevents the uptake of certain nitrogen sources when other sources are available. Inactivation of Ure2p by conversion to the prion form thus allows cells to access additional nitrogen sources [76, 69]. Jiang et al. investigated the aggregation of the full-length wild-type Ure2p and a variant which lacked residues 15-42 [77]. Data analysed in our study were extracted from figure 2D [77].

##### 2.25 WW domain

WW domain proteins are ubiquitous in eukaryotes, notable for their ultra-fast folding into anti-parallel $\beta$ sheet structures with tryptophan (W) residues at the N- and C-termini [78]. The $\lambda$ and $\varkappa$ values reported here were extracted by Knowles et al. from data in figure 2A of a study by Ferguson et al. [79, 78]. Ferguson et al. studied the aggregation of the 40-residue murine wild-type FBP28 WW domain [78].

#### 3 Data and fits

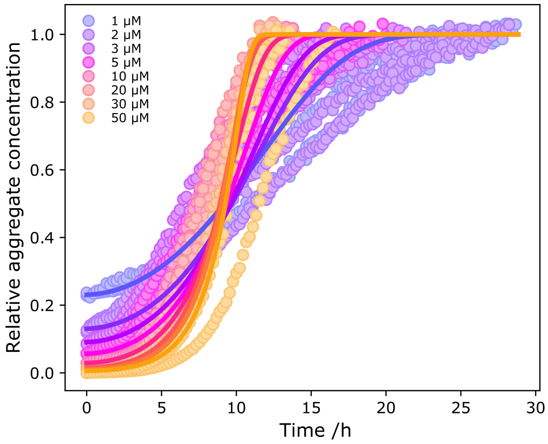


**Figure S1:** α-Synuclein aggregation kinetics. Gaspar et al. used ThT fluorescence to monitor the aggregation kinetics of different concentrations of α-synuclein monomers in the presence of 0.3 μM fibril seeds. Data were extracted from figure 2C [3].

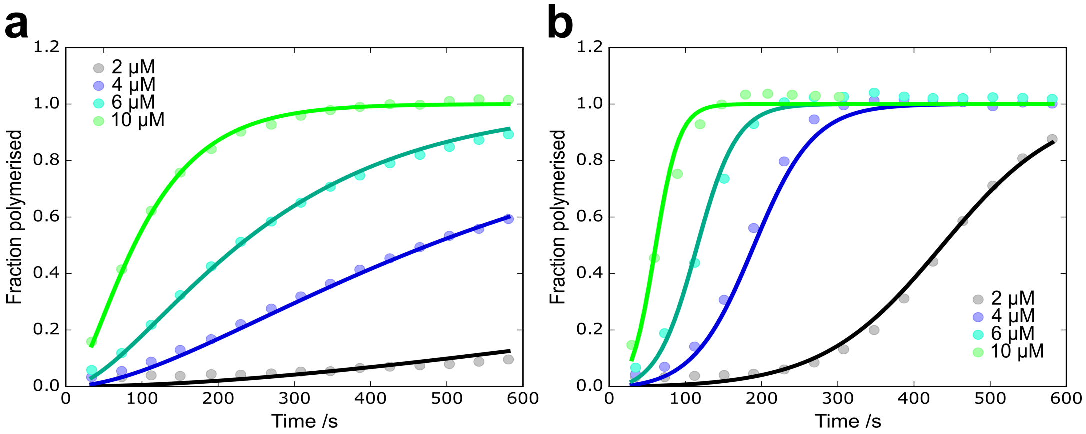


**Figure S2:** Actin polymerisation kinetics. Higgs et al. monitored the polymerisation kinetics of actin by the fluorescence of pyrene-labelled actin monomers. (a) Polymerisation kinetics of actin at different concentrations. (b) Actin polymerisation kinetics in the presence of 100 nM WA and 10 nM Arp2/3. Data in both (a) and (b) were extracted from figure 9 [14].

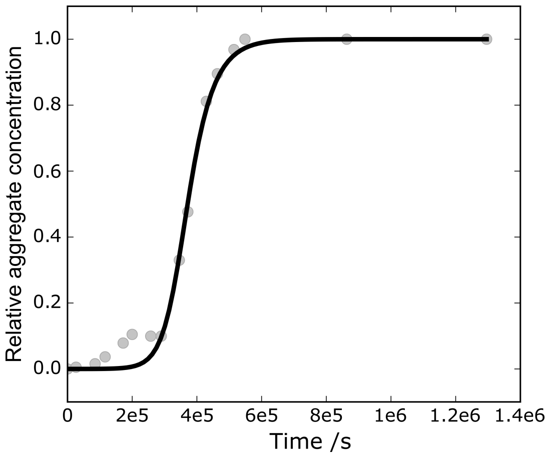


**Figure S3:** Apomyoglobin aggregation kinetics. Vilasi et al. used ThT fluorescence to monitor the aggregation kinetics of 40 µM apomyoglobin. Data were extracted from figure 3 [17].

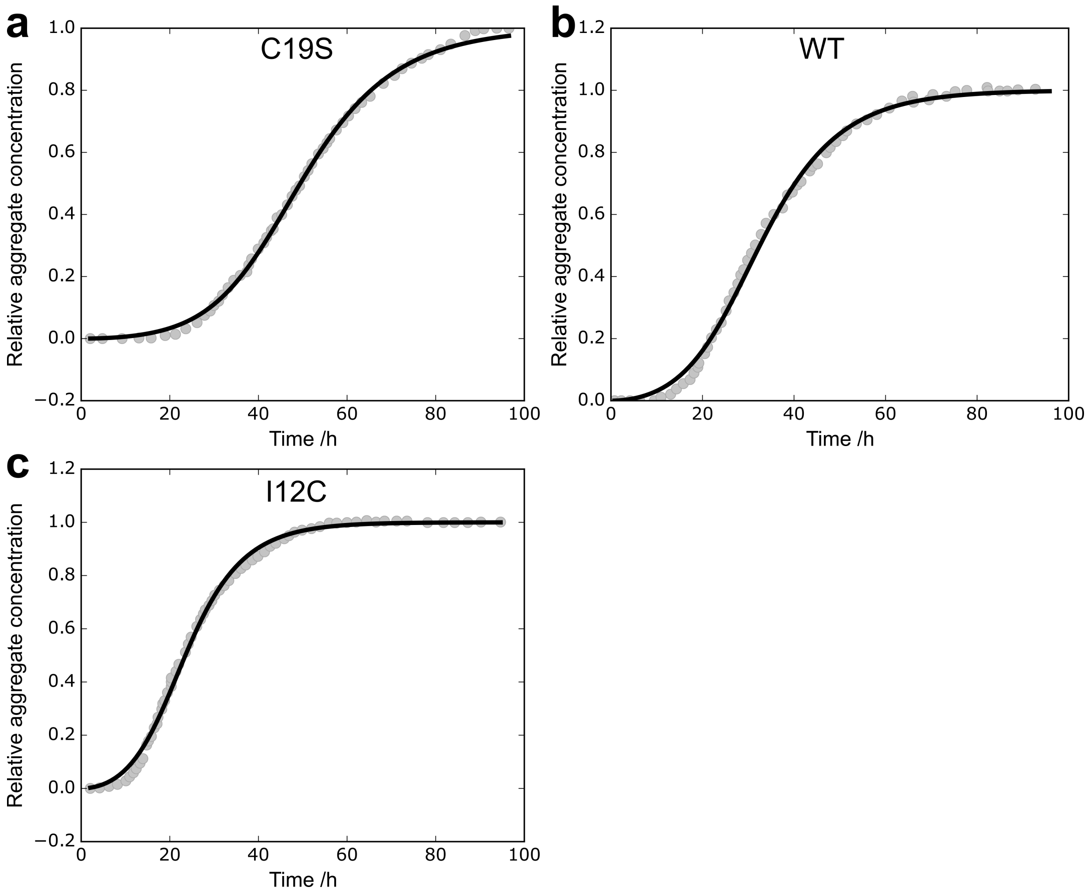


**Figure S4:** Gadm1 aggregation kinetics. Castellanos et al. used ThT fluorescence to monitor the aggregation kinetics of 173 µM WT Gadm1 and two variants. Data were extracted from figure 3a [23].

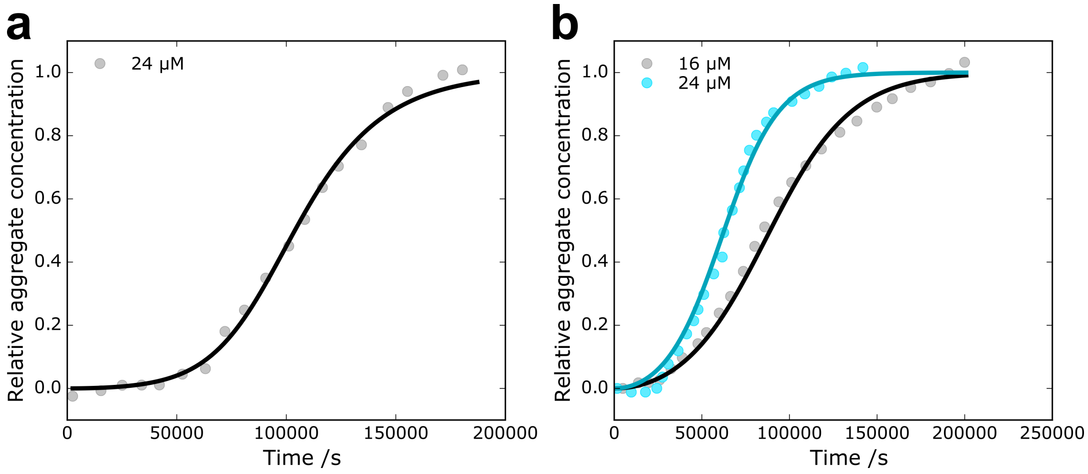


**Figure S5:** Gelsolin aggregation kinetics. Solomon et al. used ThT fluorescence to monitor the aggregation kinetics of 5 kDa (a) and 8 kDa (b) fragments of gelsolin. Data were extracted from figure 1d [24].

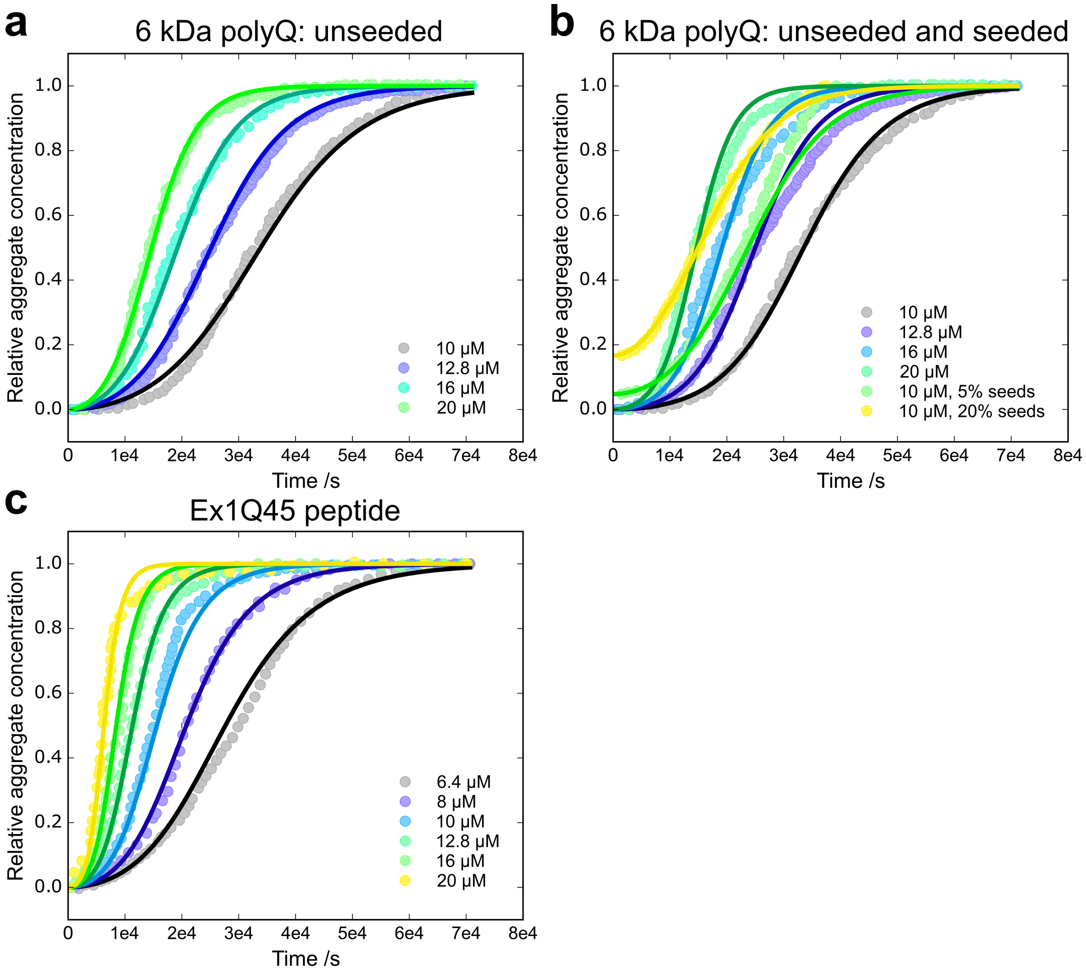


**Figure S6:** Htt exon 1/polyQ aggregation kinetics. Kakkar et al. used ThT fluorescence to monitor the aggregation kinetics of a 6 kDa peptide containing a polyQ tract of length 45 at varying concentrations in both the absence (a) and presence (b) of fibrillar seeds. They additionally studied the aggregation of a 14 kDa Huntingtin exon 1 construct, also containing a 45Q tract (c). Data in (a), (b), and (c) were extracted from figures 1a, 1a/S1b, and S1c, respectively [25].

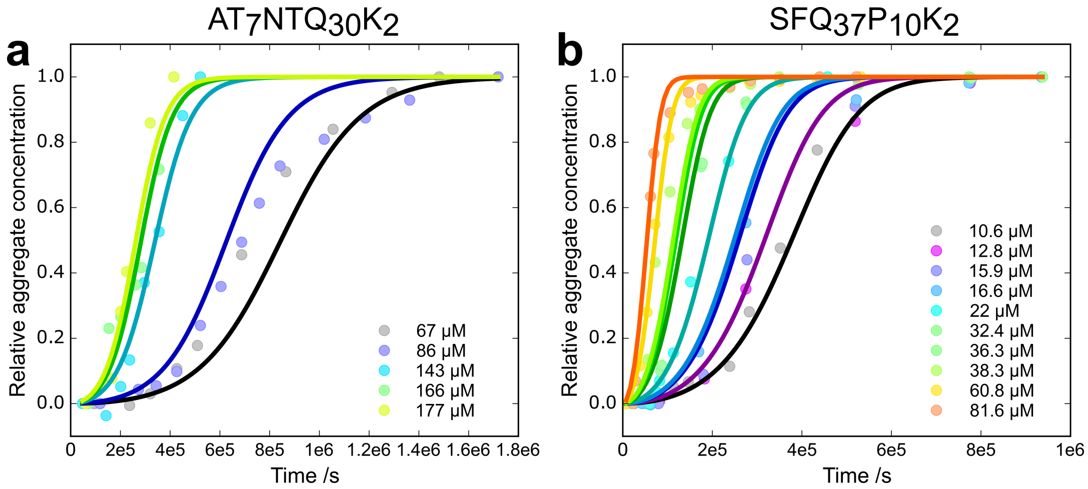


**Figure S7:** PolyQ peptide aggregation kinetics. Kar et al. monitored the aggregation kinetics of different peptides containing varying lengths of polyQ tracts, by sedimenting fibrils and quantifying the concentration of remaining monomers. We analysed their data on the concentration dependence of peptides with polyQ tracts of lengths 30 (a) and 37 (b), extracted from figures 1c and 3b, respectively [26].

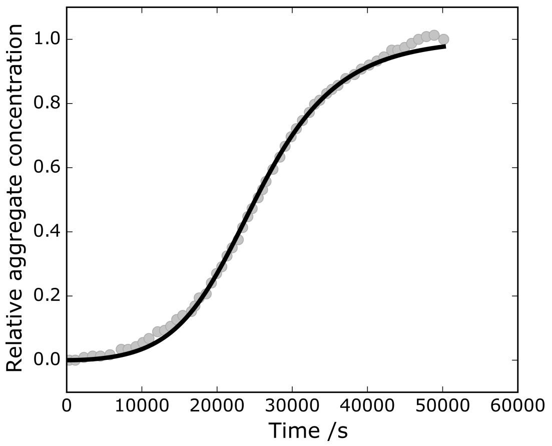


**Figure S8:** IAPP aggregation kinetics. Pilkington et al. used ThT fluorescence to monitor the aggregation kinetics of IAPP. Data were extracted from figure 1a [26].

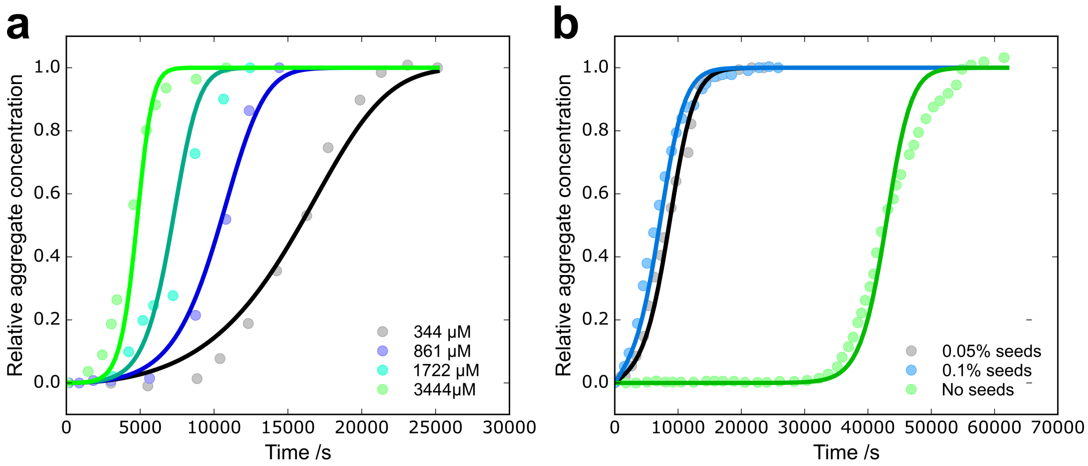


**Figure S9:** Insulin aggregation kinetics. Nielsen et al. used ThT fluorescence to monitor the concentration dependence of insulin aggregation kinetics in the absence (a) and presence (b) of seeds. In the presence of seeds, the total protein concentration was 344 μM. Data in (a) and (b) were extracted from figures 2a and 6, respectively [29].

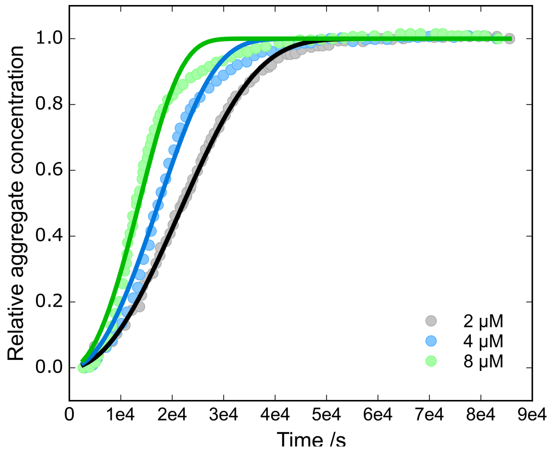


**Figure S10:** LacY aggregation kinetics. Stroobants et al. used ThT fluorescence to monitor the concentration dependence of LacY aggregation kinetics. Data were extracted from figure 3a [30].

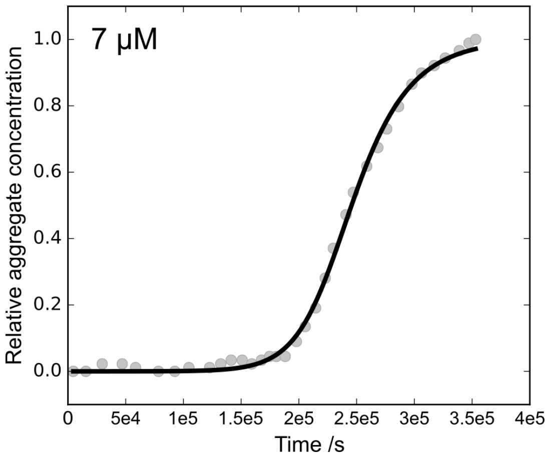


**Figure S11:** Lysozyme aggregation kinetics. Hasecke et al. used ThT fluorescence to monitor the concentration dependence of Lysozyme aggregation kinetics. At high concentrations off-path species become significantly populated, thus we use data at low concentration (7 $\mu$M) only. Data were extracted from figure 2c [31].

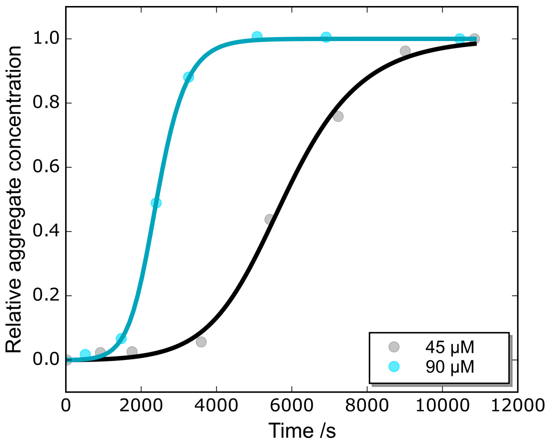


**Figure S12:** Monellin aggregation kinetics. Konno et al. used ThT fluorescence to monitor the concentration dependence of monellin aggregation kinetics. Data were extracted from figure 5a [30].

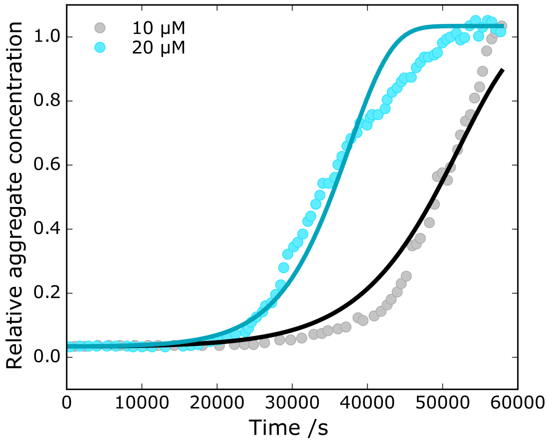


**Figure S13:** FC27 MSP2 aggregation kinetics. Adda et al. used ThT fluorescence to monitor the concentration dependence of MSP2 aggregation kinetics from the FC27 strain of *Plasmodium falciparum*. Data were extracted from figure 5 [35].

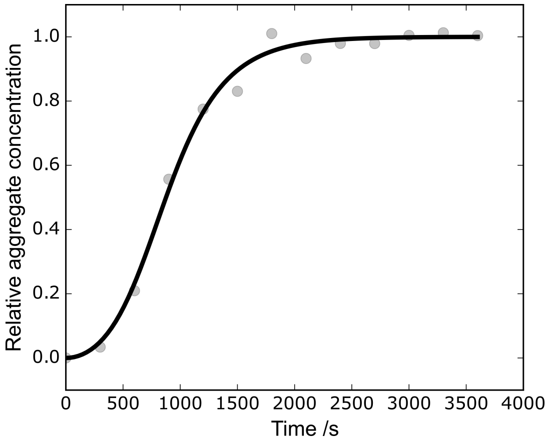


**Figure S14:** MSP(1-25)[Y7A, Y16A] aggregation kinetics. Low et al. used ThT fluorescence to monitor the aggregation of the N-terminal region of the MSP2 protein, including two mutations (Y7A and Y16A). Data were extracted from figure 6b [33].

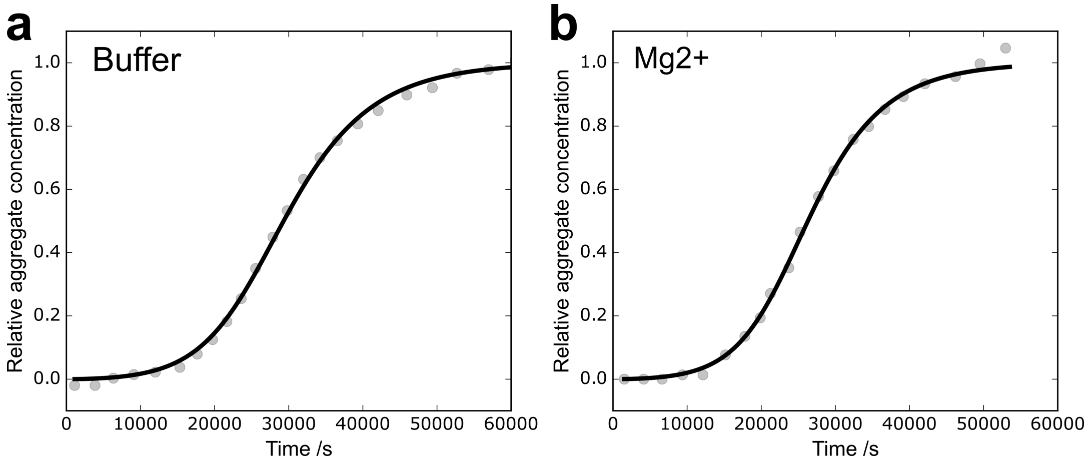


**Figure S15:** Orb2 aggregation kinetics. Bajakian et al. used ThT fluorescence to monitor the aggregation of the N-terminal region (first 87 residues) of the Orb2 protein, in the absence (a) and presence (b) of magnesium ions. The concentration of Orb2 was 17.64 µM, and the magnesium was present as a 2.5× molar excess of protein. Data were extracted from figure 6a [39].

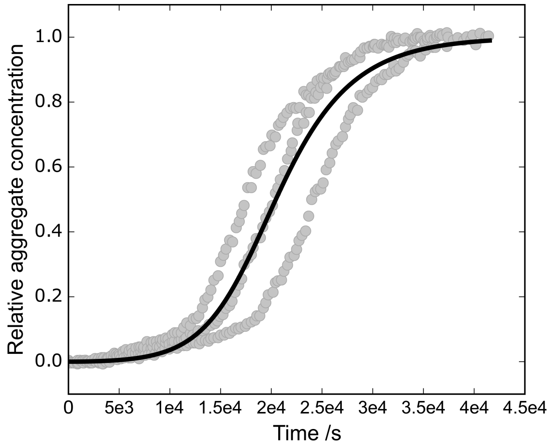


**Figure S16:** Semenogelin-derived peptide aggregation kinetics. Frohm et al. used ThT fluorescence to monitor the aggregation of a peptide composed of residues 38-48 of semenegelin at 264 µM. Data were extracted from figure 5 [55].

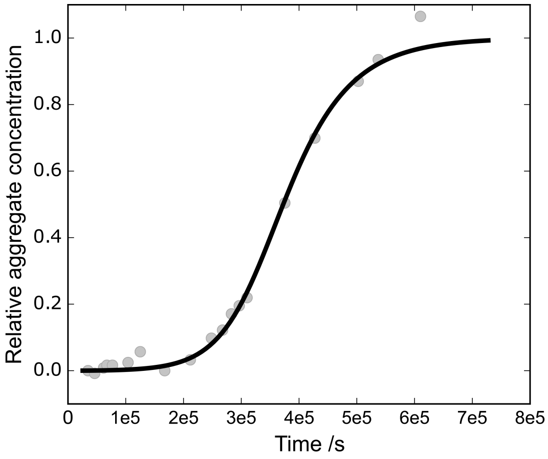


**Figure S17:** Serum amyloid (murine SAA1.1 isoform) aggregation kinetics. Srinivasan et al. used ThT fluorescence to monitor the aggregation of the murine SAA1.1 isoform of serum amyloid at a concentration of 26 µM. Data were extracted from figure 3a [59].

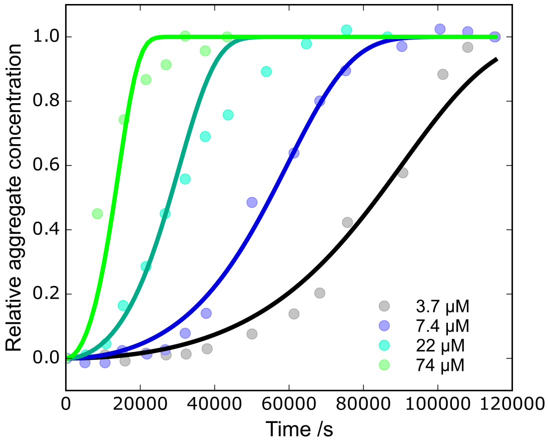


**Figure S18:** Serum amyloid (murine SAA2.2 isoform) aggregation kinetics. Ye et al. used ThT fluorescence to monitor the concentration dependent aggregation of the murine SAA2.2 isoform of serum amyloid. Data were extracted from figure 2 [62].

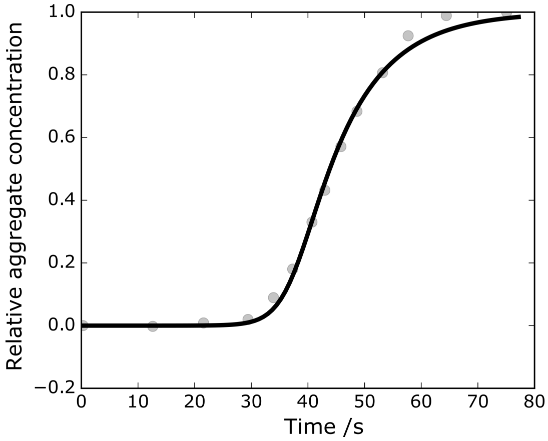


**Figure S19:** Sickle haemoglobin polymerisation kinetics. Ferrone et al. measured turbidity at 1090 nm to monitor the polymerisation of of sickle haemoglobin at 3.95 mM. In the context of sickle haemoglobin, turbidity has been shown to yield the same kinetics as NMR measurements, and can thus be related linearly to polymer concentration [80]. Data were extracted from figure 9 of Ferrone et al. [65].

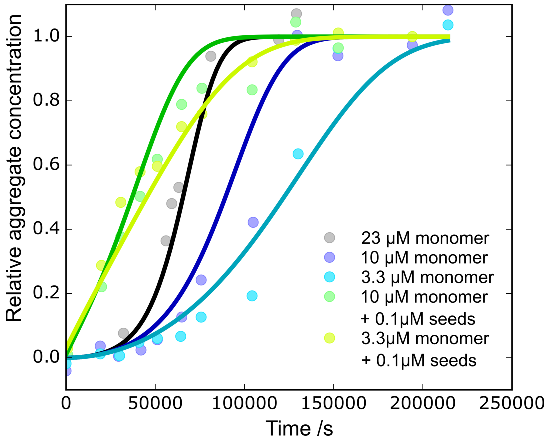


**Figure S20:** Sup35 (residues 1-253) aggregation kinetics. Glover et al. used Congo red fluorescence to monitor the aggregation of the murine SAA2.2 isoform of serum amyloid, varying concentrations of both monomers and fibril seeds. Data were extracted from figure 4 [71].

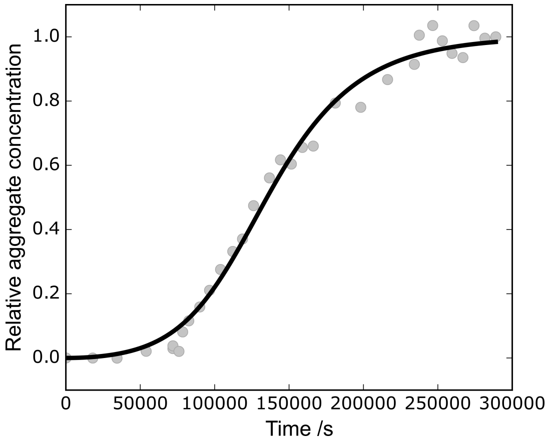


**Figure S21:** Sup35 prion-forming domain (residues 98-118) aggregation kinetics. Sant'anna et al. used ThT fluorescence to monitor the aggregation of the prion-forming domain of Sup35. Data were extracted from figure 6 [70].

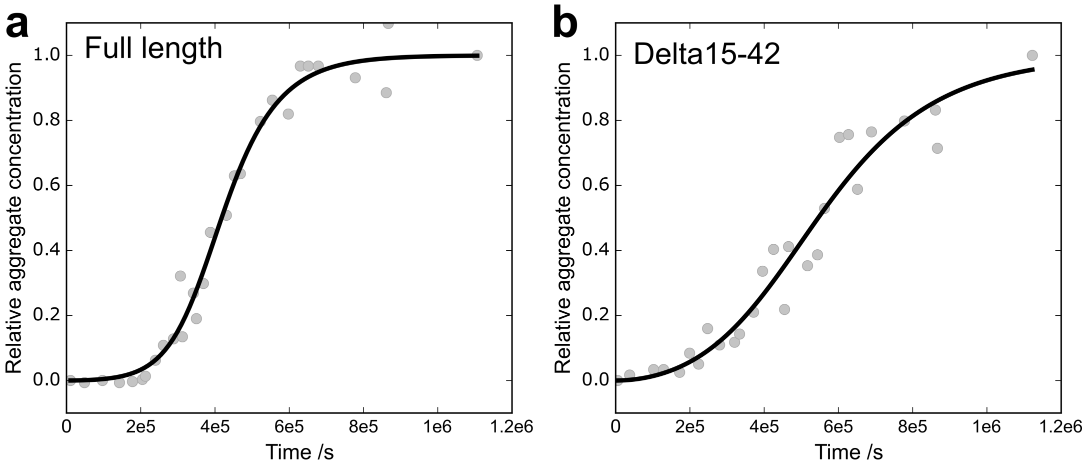


**Figure S22:** Ure2p aggregation kinetics. Jiang et al. used ThT fluorescence to monitor the aggregation of both the full length wild-type (a) and Δ15-42 variant Ure2p. Data were extracted from figure 2d [77].

| System | Source of data | *In vivo* timescale /s | | | *In vivo* concentration /M | | |
|---|---|---|---|---|---|---|---|
| | | **Minimum** | **Maximum** | **Reference(s) for timescales** | **Minimum** | **Maximum** | **Reference(s) for concentrations** |
| α-Synuclein | [3] | 6.22e7 | 6.53e8 | [81] | 2.16e-7 | 2.16e-5 | [82] |
| Aβ (all variants) | [4] | 4.04e8 | 1.06e9 | [83] | 1e-11 | 1e-5 | [84] |
| Actin | [14] | 1 | 1e3 | [85] | 5e-5 | 5e-4 | [86] |
| Actin + WAsp | [14] | 1 | 1e3 | [85] | 5e-5 | 5e-4 | [86] |
| CsgA | [20] | 3.6e3 | 3.6e5 | [20] | 1e-7 | 1e-5 | [20] |
| FapC | [20] | 3.6e3 | 3.6e5 | [20] | 1e-7 | 1e-5 | [20] |
| FC27 MSP2 | [35] | 1.8e3 | 1.44e4 | [87] | 1e-6 | 2.7e-4 | [88] |
| Gelsolin | [24] | 6.22e8 | 1.24e9 | [89] | 1.3e-8 | 2.74e-06 | [90], [91] |
| Htt | [25] | 4.67e8 | 9.64e8 | [92] | 1e-8 | 1.43e-04 | [93], [94] |
| IAPP | [28] | 3.15e7 | 3.15e8 | [95] | 1e-3 | 4e-3 | [96] |
| Prp | [49] | 7.78e6 | 5.18e7 | [97] | 1e-11 | 1e-8 | [98] |
| PSMα1 | [48] | 3.6e3 | 3.6e5 | [99] | 1e-6 | 1e-5 | [99] |
| PSMα3 | [48] | 3.6e3 | 3.6e5 | [99] | 1e-6 | 1e-5 | [99] |
| Serum amyloid | [62] | 8.29e7 | 3.40e8 | [100], [101] | 4.4e-8 | 3.7e-6 | [102], [103] |
| Sickle haemoglobin | [65] | 8.99e6 | 1.06e7 | [104], [105] | 3.93e-03 | 5.30e-03 | [106] |
| Sup35 | [71] | 5.40e3 | 1.73e5 | [107],[108] | 1e-6 | 1e-4 | [109] |
| Tau | [73], [28] | 4.04e8 | 1.06e9 | [83] | 1e-7 | 1e-5 | [73] |

**Table 1:** Biologically relevant timescales and concentrations, with the respective references, for all proteins in Fig. 3 of the main text.

| System | Data source | $\lambda$ | $\kappa$ |
|---|---|---|---|
| Apomyoglobin | Vilasi et al. [17] | 1.72e-7 | 3.06e-5 |
| Gadm1 C19S | Castellanos et al. [23] | 2.98e-6 | 3.24e-5 |
| Gadm1 I12C | Castellanos et al. [23] | 9.72e-6 | 5.16e-5 |
| Gadm1 WT | Castellanos et al. [23] | 6.67e-6 | 3.88e-5 |
| Gelsolin 5 kDa fragment | Solomon et al. [24] | 4.05e-6 | 6.21e-5 |
| IAPP | Pilkington et al. [27] | 2.21e-5 | 2.25e-4 |
| Lysozyme | Hasecke et al. [31] | 1.87e-7 | 4.94e-5 |
| Orb2 without $Mg^{2+}$ | Bajakian et al. [39] | 1.57e-5 | 2.12e-4 |
| Orb2 with $Mg^{2+}$ | Bajakian et al. [39] | 1.61e-5 | 2.45e-4 |
| Semenogelin | Frohm et al. [55] | 1.80e-5 | 3.34e-4 |
| Serum amyloid (Saa1.1) | Srinivasan et al. [59] | 6.26e-7 | 2.17e-5 |
| Sickle haemoglobin | Ferrone et al. [65] | 1.5e-4 | 4.1e-1 |
| Sup35 | Santanna et al. [70] | 4.32e-6 | 4.05e-5 |
| Ure2p $\Delta$15-42 | Jiang et al. [77] | 1.65e-6 | 7.44e-6 |
| Ure2p WT | Jiang et al. [77] | 8.73e-7 | 1.64e-5 |

**Table 2:** Proteins shown in the main text, where no reaction order could be determined. Proteins for which the reaction order could be determined are given in table 3 below. Values for some additional variants of the same proteins that were not shown in the main text Figure for lack of space are included here.

| System | Data source | $k_+k_n$ in $\mathbf{M}^{-n_c}\mathbf{s}{-2}$ | $k_+k_2$ in $\mathbf{M}^{-n_2-1}\mathbf{s}{-2}$ | $n_c$ | $n_2$ | Concentration used in Fig.2 / M | Comment |
|---|---|---|---|---|---|---|---|
| α-Synuclein | [3] | † | 1.1e-3 | † | 0 | 5e-5 | $K_E$ = 1.4e-5 M |
| Aβ40 | [5] | 0.31 | 1.0e9 | 2 | 2 | 3e-6 | $K_M$ = 3.6e-11 $M^2$ |
| Aβ42 | [6] | 4.8e1 | 8.6e10 | 2 | 2 | 3e-6 | $K_M$ = 9.2e-12 $M^2$ |
| Aβ42 (fragmentation) | [9] | n/a | 0.2 | n/a | 0 | n/a | |
| Aβ42 (D23N) | [6] | 3.8e2 | 1.1e13 | 2 | 2 | 3e-6 | $K_M$ = 9.0e-14 $M^2$ |
| Aβ42 (E22G) | [6] | 4.2e3 | 8.1e12 | 2 | 2 | 3e-6 | $K_M$ = 2.4e-13 $M^2$ |
| Aβ42 (E22Q) | [6] | 4.1e2 | 5.5e11 | 2 | 2 | 3e-6 | $K_M$ = 6.4e-13 $M^2$ |
| Aβ42 (V18S+A21S) | [8] | 2.6e1 | 1.6e10 | 2 | 2 | 3e-6 | $K_M$ = 1.6e-12 $M^2$ |
| Actin | [14] | 3.8e14 | † | 3.7 | † | 2e-6 | |
| Actin + WAsp | [14] | 3.0e11 | 1.4e7 | 3.1 | 1.0 | 2e-6 | |
| CsgA | [20] | 8.5e-3 | † | 1.2 | † | 4e-6 | |
| FapC | [20] | 1.4e-4 | † | 1.3 | † | 1e-4 | |
| FC27 MSP2 | [35] | 1.88e-7 | 9.7e-4 | 0.8 | 0 | 1e-5 | |
| Gelsolin 8 kDa fragment | [24] | 4.8e-4 | 3.1 | 1.5 | 1.0 | 1.6e-5 | |
| Htt Ex45 | [25] | 12 | 4.5e7 | 2 | 2 | 1.28e-5 | |
| Htt 6 kDa polyQ | [25] | 1.1e3 | 2.3e3 | 2.5 | 1.3 | 1.28e-5 | |
| Htt Q30 | [26] | 0.5 | 5.8e-3 | 3.0 | 1.0 | 8.6e-5 | |
| Htt Q37 | [26] | 0.3 | 1.6e-3 | 2.3 | 0.5 | 3.24e-5 | |
| IAPP | [28] | 1.7e5 | 1.1e7 | 3 | 1.6 | 5e-6 | |
| Insulin | [29] | 1.0e-9 | 1.6e-2 | 0 | 0.7 | 3.44e-4 | |
| LacY | [30] | 7.2e-6 | 8.8e-4 | 0.7 | 0 | 4e-6 | |
| Monellin | [32] | 0.2 | 8.8e5 | 1.8 | 1.8 | 4.5e-5 | |
| PrP | [49] | † | 6.0e-2 | † | 0 | 1e-6 | |
| Psmα1 | [48] | 5.4e-12 | 1.0e-5 | 0 | 0 | 1.11e-4 | |
| Psmα3 | [48] | 2.1e-5 | 0.4 | 0.6 | 0.1 | 1.53e-4 | |
| Serum amyloid | [62] | 1.4e-3 | 2.7e-4 | 1.4 | 0 | 2.2e-5 | |
| Sup35 | [71] | 3.0e-11 | 4.3e-3 | 0.3 | 0 | 1e-5 | |
| Tau (WT) | [73] | n/a | 2.4e-7 | n/a | 0 | 2e-6 | |
| Tau (304-380) | [72] | 1e-12 | 6e11 | 0 | 2 | 3.2e-6 | $K_M$ = 3e-8 $M^2$ |

**Table 3:** Proteins shown in the main text, where a reaction order could be determined. When experiments were seeded or self-replication processes were too slow to be detected this is denoted by †. In those cases, upper bounds for $\lambda$ or $\kappa$, were used instead, calculated as outlined in the Methods section. "A$\beta$42 (fragmentation)" simply records the estimate of the rate of fragmentation A$\beta$42 (see section on A$\beta$). The tau datasets from Kundel et al.[73] use heparin to artificially induce primary nucleation, hence the primary rate constants and reaction orders observed in that experiment are not used here (see section on tau). Some datasets had to be analysed with a model including saturation of secondary nucleation or elongation, the respective values for the saturation constants, $K_M$ and $K_E$, respectively, are given in the "comments" column. For details on the models please refer to the cited papers as well as Meisl et al. [1].

### 4 Investigation of structure-mechanism correlation

In order to investigate whether any specific structural features could be identified that correlate with the propensity for self-replication to dominate over de novo formation, we used a recently developed method to identify sequence patterns, see Fig. S23. This analysis yields 36 different types of patterning score, for each of which we then computed the Pearson correlation coefficient, and its associated p-value for non-correlation, with our measure for dominance of self-replication, $log(\kappa/\lambda)$. A weak correlation of $log(\kappa/\lambda)$ with some patterning scores is observed, however, when the Bonferroni correction is applied to account for the fact that there are 36 different scores to correlate to, none of these correlations remain significant ($p > 0.05$). While there are likely specific structural features that promote or prevent self-replication, our current dataset is likely not extensive enough to provide statistically significant insights and considerable additional work will be needed to elucidate these features.

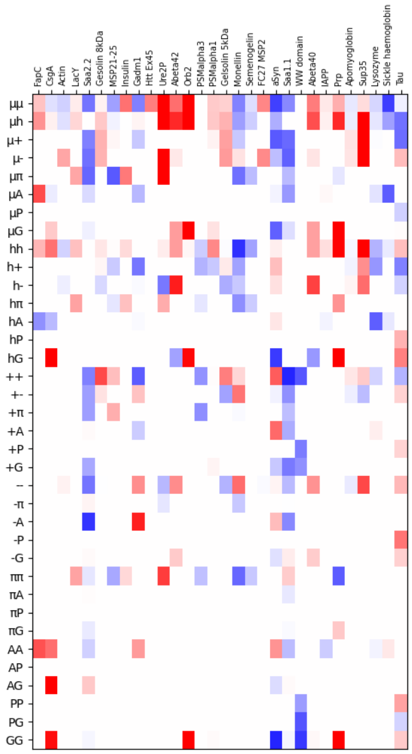


**Figure S23: Z-scores from NARDINI analysis[110].** Each column is a different protein (when data from multiple forms of the protein were available, the WT was chosen), each row a different patterning parameter, the colour denotes the z-score, which quantifies how much each patterning score differs from a random sequence, with blue and red indicating more and less uniform dispersion, respectively, than would be expected in a scrambled sequence. The data are ordered by increasing $\kappa/\lambda$ ratio from left to right, i.e. proteins for which de novo formation dominates are on the left, those for which self-replication dominates are on the right.

[3] Ricardo Gaspar, Georg Meisl, Alexander K. Buell, Laurence Young, Clemens F.

Kaminski, Tuomas P. J. Knowles, Emma Sparr, and Sara Linse. Secondary nucleation of monomers on fibril surface dominates $\alpha$-synuclein aggregation and provides autocatalytic amyloid amplification. *Quarterly Reviews of Biophysics*, 50:E6, 2017.

[28] Diana C. Rodriguez Camargo, Sean Chia, Joseph Menzies, Benedetta Mannini, Georg Meisl, Martin Lundqvist, Christin Pohl, Katja Bernfur, Veronica Lattanzi, Johnny Habchi, Samuel IA Cohen, Tuomas P. J Knowles, Michele Vendruscolo, and Sara Linse. Surface-catalyzed secondary nucleation dominates the generation of toxic iapp aggregates. *Frontiers in Molecular Biosciences*, 8, 2021.

[29] L. Nielsen, R. Khurana, A. Coats, S. Frokjaer, J. Brange, S. Vyas, V. N. Uversky, and A. L. Fink. Effect of environmental factors on the kinetics of insulin fibril formation: elucidation of the molecular mechanism. *Biochemistry*, 40(20):6036–6046, May 2001.

[30] Karen Stroobants, Janet R. Kumita, Nicola J. Harris, Dimitri Y. Chirgadze, Christopher M. Dobson, Paula J. Booth, and Michele Vendruscolo. Amyloid-like

fibrils from an α-helical transmembrane protein. *Biochemistry*, 56(25):3225–3233, 2017.

[86] H Lodish, A Berk, and SL Zipursky. *Molecular Cell Biology, 4th edition. Section 18.2, The Dynamics of Actin Assembly.* New York: W. H. Freeman, 2000.

[95] Vivian A. Fonseca. Defining and characterizing the progression of type 2 diabetes. *Diabetes Care*, 32(suppl 2):S151–S156, 2009.

[96] Emma T. A. S. Jaikaran, Melanie R. Nilsson, and Anne Clark. Pancreatic beta-cell granule peptides form heteromolecular complexes which inhibit islet amyloid polypeptide fibril formation. *Biochemical Journal*, 377(3):709–716, 02 2004.

[97] M. Pocchiari, M. Puopolo, E. A. Croes, H. Budka, E. Gelpi, S. Collins, V. Lewis, T. Sutcliffe, A. Guilivi, N. Delasnerie-Laupretre, J.-P. Brandel, A. Alperovitch, I. Zerr, S. Poser, H. A. Kretzschmar, A. Ladogana, I. Rietvald, E. Mitrova, P. Martinez-Martin, J. de Pedro-Cuesta, M. Glatzel, A. Aguzzi, S. Cooper, J. Mackenzie, C. M. van Duijn, and R. G. Will. Predictors of survival in sporadic creutzfeldt–jakob disease and other human transmissible spongiform encephalopathies. *Brain*, 127(10):2348–2359, 09 2004.

[98] Sonia M. Vallabh, Chloe K. Nobuhara, Franc Llorens, Inga Zerr, Piero Parchi, Sabina Capellari, Eric Kuhn, Jacob Klickstein, Jiri G. Safar, Flavia C. Nery,

Kathryn J. Swoboda, Michael D. Geschwind, Henrik Zetterberg, Steven E. Arnold, Eric Vallabh Minikel, and Stuart L. Schreiber. Prion protein quantification in human cerebrospinal fluid as a tool for prion disease drug development. *Proceedings of the National Academy of Sciences*, 116(16):7793–7798, 2019.

[106] PR Sarma. *Red Cell Indices. In: Walker HK, Hall WD, Hurst JW, editors. Clinical Methods: The History, Physical, and Laboratory Examinations. 3rd edition.* Boston: Butterworths, 1990.

[107] Nadège Minois, Magdalena Frajnt, Chris Wilson, and James W. Vaupel. Advances in measuring lifespan in the yeast saccharomyces cerevisiae. *Proceedings of the National Academy of Sciences*, 102(2):402–406, 2005.

[108] Libuše Váchová, Michal Čáp, and Zdena Palková. Yeast colonies: A model for studies of aging, environmental adaptation, and longevity. *Oxidative Medicine and Cellular Longevity*, 2012:601836, Aug 2012.

[109] Tarique Khan, Tejbir S. Kandola, Jianzheng Wu, Shriram Venkatesan, Ellen Ketter, Jeffrey J. Lange, Alejandro Rodríguez Gama, Andrew Box, Jay R. Unruh, Malcolm Cook, and Randal Halfmann. Quantifying nucleation in vivo reveals the physical basis of prion-like phase behavior. *Molecular Cell*, 71(1):155–168.e7, 2018.

[110] Megan C. Cohan, Min Kyung Shinn, Jared M. Lalmansingh, and Rohit V. Pappu. Uncovering non-random binary patterns within sequences of intrinsically disordered proteins. *Journal of Molecular Biology*, 434(2):167373, 2022.
